# Dual targeting of PDPK1 and BRAF V600E is synthetically lethal

**DOI:** 10.64898/2026.03.15.711663

**Authors:** Tejinder Pal Khaket, Chandrayee Ghosh, Zhongyue Yang, Myriem Boufraqech, Jiangnan Hu, Eden D. Alamaw, Thorkell Andresson, Maura O’Neill, Ya-Qin Zhang, Min Shen, Bereketeab Haileselassie, Electron Kebebew

## Abstract

PDPK1 functions downstream of PI3K and is essential for activating AKT and other AGC kinases. Although PDPK1 has a central role in the PI3K/AKT/mTOR signaling pathway, there has been limited evaluation of it as a target for cancer therapy. Anaplastic thyroid cancer (ATC) has one of the highest mortality rates of all human malignancies. Although combined BRAF and MEK inhibition in BRAF V600E mutant ATC (present in 45% of cases) results in response, resistance is common, and there is no curative treatment for ATC. The majority (up to 95.8%) of ATC cases have activation in the PI3K/AKT/mTOR and RAS/RAF/MEK/MAPK pathways due to genetic alterations involved in these pathways. In this study, we investigated PDPK1 as a therapeutic target for ATC. We used in vitro, ex vivo, and in vivo ATC models to evaluate the effect of targeting PDPK1 (BX795) alone and in combination with BRAF V600E inhibition (dabrafenib), and the mechanism of action that resulted in ATC cell death. BX795 monotherapy significantly reduced ATC cell proliferation, invasion, colony formation, and spheroid size. Combination BX795 and dabrafenib treatment had strong synergistic anticancer activity in BRAF V600E-mutant ATC models and led to simultaneous and sustained suppression of PDPK1/AKT and MAPK signaling, preventing the compensatory pathway reactivation observed with single-agent treatment. Mechanistically, combined inhibition induced pronounced oxidative stress, DNA damage, and G2-phase cell-cycle arrest, accompanied by mitochondrial dysfunction and robust activation of apoptosis in ATC cells. These effects resulted in marked tumor regression in in vitro, ex vivo, and in vivo ATC models. Our findings identify PDPK1 as a critical therapeutic vulnerability in ATC. Co-targeting PDPK1 and BRAF V600E produces potent synergistic anticancer activity by shutting down convergent oncogenic signaling pathways and amplifying apoptotic stress responses. These data support PDPK1 inhibition alone and in combination with BRAF blockade as a promising therapeutic strategy in BRAF V600E-mutant cancers.

## INTRODUCTION

Anaplastic thyroid cancer (ATC) is a rare (incidence of 1–2 cases per million people) and lethal cancer. Although ATC accounts for 1%–2% of all thyroid cancer cases, it is responsible for two-thirds of all thyroid cancer deaths and has a median survival time of only 6 months (1–3). Recent advances in the understanding of genetic alterations in ATC have revealed that the majority (up to 95.8%) of ATC cases have activation in the PI3K/AKT/mTOR and MAPK/ERK (also known as RAS/RAF/MEK/MAPK) pathways due to genomic alterations (driver mutations and genomic gains/losses) involved in these pathways (4,5). Targeting driver mutations, such as *BRAF* V600E (present in 45% of ATC cases), in mutant ATC elicits a response in some cases, but development of resistance to treatment and treatment-related toxicity that results in treatment stoppage are common (3,6–10). Thus, there is an urgent need to develop new therapeutic strategies for ATC that elicit a significant and durable response and thereby extend patient survival.

Although inhibition of the MAPK/ERK pathway in *BRAF* V600E–mutant ATC, using BRAF and MEK inhibitors, is effective, compensatory upregulation and acquired mutations in the PI3K/AKT/mTOR pathway commonly undermine this strategy, leading to therapeutic resistance (11–16). Within the PI3K/AKT/mTOR signaling pathway, 3-phosphoinositide-dependent kinase 1 (PDPK1 or PDK1) functions downstream of PI3K and is essential for activating AKT and other AGC kinases (17,18). Although PDPK1 has a central role in the PI3K/AKT/mTOR signaling pathway, there has been limited investigation of it as a target for cancer therapy (18). Gene mutations that lead to activation of the PI3K/AKT/mTOR pathway are common in ATC and occur in genes upstream of PDPK1. However, direct evidence of PDPK1 phosphorylation in ATC has not yet been reported, representing an important knowledge gap. Phosphorylation of PDPK1 at Ser241 is essential for its activity in phosphorylating AKT at Thr308 and its association with cancer initiation and progression (19). PDPK1 overexpression drives cancer initiation and progression in a range of malignancies, and is associated with more aggressive tumors (20–38). PDPK1 is also overexpressed in thyroid cancer, where higher levels are associated with more advanced disease and ATC (39). Accumulating evidence indicates that PDPK1-mediated AKT activation plays a central role in cancer initiation and progression (40–44).

Investigating PDPK1 activation in ATC could provide both mechanistic insight and define the potential of targeting it for ATC therapy and other cancers with activation in the PI3K/AKT/mTOR pathway. Therefore, we evaluated the anticancer activity of targeting PDPK1 using *in vitro*, *ex vivo*, and *in vivo* models in ATC with and without *BRAF* V600E inhibition.

## MATERIALS AND METHODS

### Materials

The materials used in this study are listed in Tables S1 (list of antibodies), S2 (list of chemicals), and S3 (cell lines and their mutation status).

### Cell lines and culture conditions

The homozygous mutated *BRAF* V600E 8505C, FTC133 (homozygous mutated *PTEN*), and FTC236 (homozygous mutated *PTEN*) human thyroid cancer cell lines were purchased from the European Collection of Cell Culture (Salisbury, UK). The heterozygous mutated *BRAF* V600E human ATC cell line SW1736 and the *BRAF* wild type ATC cell line C643 (heterozygous mutated *PTEN*) were purchased from Cell Lines Service GmbH (Eppelheim, Germany). The THJ-11T (*KRAS* mutant), THJ-29T, and THJ-16T (*PIK3CA* mutant) ATC cell lines were a kind gift from Dr. John A. Copland III (Mayo Clinic, Jacksonville, FL). The BCPAP (homozygous mutated *BRAF* V600E) cell line was purchased from Leibniz Institute DSMZ (Lower Saxony, Germany). The TPC1 (*RET/PTC1* rearrangement) cell line was purchased from Sigma-Aldrich, Inc. (St. Louis, MO, USA). Details of cell line–specific mutations are provided in Table S3. The HEK293 cell line was purchased from American Type Culture Collection (Manassas, VA, USA). All cells were cultured in Dulbecco’s Modified Eagle’s Medium (DMEM, which contains 4,500 mg/L of D-glucose, 2 mmol/L of L-glutamine, and 110 mg/L of sodium pyruvate) supplemented with 10% fetal bovine serum (FBS), and antibiotics (penicillin and streptomycin at 10,000 U/mL) at 37 °C and 5% CO_2_. Two patient-derived *BRAF* V600E–mutant ATC cell lines—ATC-01, derived from a treatment-naïve patient’s primary *BRAF* V600E–mutant tumor, and ATC-02, derived from a *BRAF* V600E–mutant residual tumor following combination therapy with dabrafenib (Dab) and trametinib (*BRAF* and MEK inhibitors, respectively)—were maintained in DMEM:F12 supplemented with 15% FBS and antibiotics (penicillin and streptomycin at 10,000 U/mL) at 37 °C and 5% CO_2_ in a humidified incubator.

### Quantitative high-throughput screening

Quantitative high-throughput screening was used to screen the 8505C, THJ-11T, and THJ-16T ATC cell lines based on the National Center for Advancing Translational Sciences (NCATS) Pharmaceutical Collection (NPC) and the Mechanism Interrogation PlatE (MIPE) library with 4,991 small-molecule drugs and investigational compounds. Cell viability was measured using a luciferase-coupled adenosine triphosphate (ATP) quantitation assay (CellTiter-Glo®, Promega, Madison, WI, USA). To identify the optimal PDPK1 inhibitor, drug response data of PDPK1 inhibitors (BX-912, DTXSID90678515, OSU-03012, GSK-2334470, and BX795) present in the high-throughput screening library were analyzed.

### Cell proliferation assay

Cells were seeded in 96-well black plates (Greiner Bio-One, Kremsmünster, Austria) at a density of 1.5 × 10^3^ cells per well. After 24 h, cells were treated in triplicate with the indicated concentrations of Dab, BX795, and other small molecules for 48 h at 37 °C and 5% CO_2_ in a humidified incubator. Dimethyl sulfoxide (DMSO) was used as a vehicle control. CyQUANT cell proliferation assays were performed according to the manufacturer’s instructions, and the results were analyzed using a SpectraMax i3x Multi-Mode Microplate Reader (Molecular Devices, San Jose, CA, USA) at an excitation wavelength of 480 nm and an emission wavelength of 530 nm. The relative fold change was measured as compared to the control. Statistical analyses were performed using GraphPad Prism version 10 (GraphPad Software Inc., San Diego, CA, USA).

### Calculation of combination drug activity

After treating ATC cells with various concentrations of DMSO (control), BX795, Dab, or their combination (BX795 + Dab) for 48 h, cell proliferation was measured. Then, drug responses were analyzed using the CompuSyn software (Version 1.0) based on the Chou–Talalay method (45) to determine synergy, additivity, or antagonism. The combination index (CI) was calculated, where CI < 1 indicates synergy, CI = 1 indicates an additive effect, and CI > 1 indicates antagonism. The values were derived from drug combinations applied at non-constant ratios.

### Clonogenic assay

Cells (1 × 10^3^ cells/well) were seeded in 12-well plates and allowed to adhere overnight. Then, cells were treated in triplicate with DMSO (control), BX795 (2.5 µM), Dab (2.5 µM), or their combination (BX795 + Dab) for 48 h, after which time the medium was replaced with fresh complete DMEM, and colonies were allowed to form for up to 8–12 days. Colonies were fixed with 4% buffered paraformaldehyde for 10 min at room temperature, washed with phosphate-buffered saline (PBS), and then stained with 0.5% crystal violet for 30 min at room temperature with continuous shaking. Finally, stained colonies were washed with water, and the ImageJ software (National Institutes of Health, Bethesda, MD, USA) was used for quantification. GraphPad Prism version 10 was used for statistical analysis.

### Migration assay

A wound-healing assay was performed to analyze cellular migration. In brief, a scratch (wound) was generated using a sterile pipette tip, and cells were washed with PBS; treated in triplicate with DMSO (control), BX795 (2.5 µM), Dab (2.5 µM), or their combination (BX795 + Dab); and incubated in DMEM supplemented with 2% FBS for 16–24 h. Finally, wound-healing was measured using ImageJ. The results are presented as the relative change in cellular migration compared to the control.

### Spheroid formation assay

Mutant *BRAF* V600E ATC cells (8505C, SW1736, ATC01, and ATC02) were seeded in round bottom, ultra-low attachment 96-well plates (1.5 × 10^3^/well) using 3D Tumorsphere Medium XF (PromoCell, Heidelberg, Germany) and placed in a humidified incubator at 37 °C with 5% CO_2_ for spheroid formation. After 72 h, spheroids were treated with DMSO (control), BX795 (2.5 µM), Dab (2.5 µM), or their combination (BX795 + Dab), and monitored over 7 days. Images of each spheroid were acquired after 0 and 8 days of treatment. The size was quantified using ImageJ. The data are presented as the relative change in spheroid size compared to the control.

### Quantitative proteomic and phosphoproteomic analysis

#### Sample preparation and tandem mass tag (TMT) labeling

Mutant BRAF V600E ATC cell lines were treated with DMSO (control), BX795 (2.5 µM), Dab (2.5 µM), or their combination (BX795 + Dab) for 48 h. Cell pellets were resuspended in 200 µL of EasyPep™ lysis buffer (Thermo Fisher Scientific, Waltham, MA, USA) supplemented with nuclease (1 µL), protease inhibitors, and PhosSTOP (Sigma-Aldrich, Inc., St. Louis, MO, USA). Protein concentrations were determined with the BCA protein assay, and 100 µg of protein per sample was mixed with 50 µL of reduction reagent and 50 µL of alkylation reagent. Samples were digested overnight at 37 °C with 6 µg of trypsin/Lys-C at 750 rpm. The following day, 200 µg of the TMTpro reagent (Thermo Scientific) was added, and samples were incubated for 1 h at 25 °C. Labeling reactions were quenched with 5% hydroxylamine and 20% formic acid (50 µL, 10 min). TMT-labeled peptides were combined, cleaned using the EasyPep™ Maxi Kit, and dried in a vacuum concentrator.

#### Phosphopeptide enrichment

Sequential phosphopeptide enrichment was performed using the High-Select TiO_2_ and Fe-NTA IMAC kits (Thermo Fisher Scientific) following the manufacturer’s protocols. Peptides were first enriched by TiO_2_; the TiO_2_ flow-through and wash fractions were collected, dried, and subjected to a second enrichment using immobilize metal affinity chromatography (IMAC). Eluted phosphopeptides from both steps were combined, dried, and reserved for liquid chromatography–tandem mass spectrometry (LC–MS/MS) analysis. IMAC flow-through and wash fractions were saved for high-pH fractionation.

#### High-pH reverse-phase fractionation

IMAC flow-through peptides were resuspended in 100 µL of LC/MS-grade water, and 85 µL was fractionated by high-pH reverse-phase ultra-performance liquid chromatography (UPLC) (ACQUITY, Waters, Milford, MA). Peptides were separated on a 150 mm × 3.0 mm XBridge Peptide BEH C18 column (2.5 µm) at 0.35 mL/min. After a 5-min wash with mobile phase A (10 mM ammonium formate, pH 9.4), peptides were eluted using a gradient of 10%–50% mobile phase B (10 mM ammonium formate/90% acetonitrile, pH 9.6; 5–60 min), followed by 50%–75% B (60–70 min). Fractions were collected every minute, generating 60 fractions, which were pooled into 12 final fractions, dried, and stored at −80 °C until analysis.

#### LC–MS/MS analysis

Dried fractions were resuspended in 0.1% formic acid, and 5 µL was injected into a Dionex UltiMate 3000 RSLC system coupled to an Orbitrap Eclipse mass spectrometer (Thermo Fisher Scientific) equipped with FAIMS and an EasySpray source. Chromatographic separation was performed at 300 µL/min using solvent A (0.1% formic acid) and solvent B (80% acetonitrile/0.1% formic acid) with the following gradient: 5%–7% B (1 min), 7%–30% B (83 min), 30–59% B (25 min), 50%–95% B (4 min), 95% B (7 min), and re-equilibration at 5% B (17 min). The mass spectrometer was operated in TopSpeed mode using four FAIMS compensation voltages (−45, −55, −65, and −75). MS1 scans were acquired in the Orbitrap at 120,000 resolutions with an automatic gain control (AGC) target of 4 × 10^5^ and a maximum injection time of 50 ms over 375–1600 m/z. MS2 spectra were acquired using the TurboTMT method at 15,000 resolutions (an AGC target of 5 × 10^4^, an isolation window of 1.6 m/z, a higher-energy collisional dissociation [HCD] of 30%, and a minimum intensity of 2.5 × 10^4^). Monoisotopic precursor selection, Easy-IC internal calibration, and advanced peak determination were enabled. For phosphopeptide runs, two FAIMS methods were used, each containing four compensation voltages (CVs) (a cycle time of 0.75 s per CV; 3 s total): Method 1 (−45, −55, −65, and −75) and Method 2 (−50, −60, −70, and −80).

#### Database searching and quantification

Raw files were processed in Proteome Discoverer using the Sequest HT node. Spectra were searched against the UniProt human database with full trypsin specificity, allowing up to two missed cleavages and a minimum peptide length of six residues. Tolerance was set to 10 ppm for MS1 and 0.02 Da for MS2. Carbamidomethylation of cysteine, TMTpro labeling of lysine and peptide N-termini, and phosphorylation of serine/threonine/tyrosine were specified as static or variable modifications as appropriate. Methionine oxidation was included as a variable modification. PSM-, peptide-, and protein-level false-discovery rates (FDRs) were controlled with Percolator. Reporter ion intensities were quantified using the Reporter Ion Quantifier node and normalized to the total peptide abundance per channel.

#### Proteomic and phosphoproteomic data analysis and pathway enrichment

All proteomic and phosphoproteomic analyses were performed in R version 4.2 using a reproducible workflow. The data were imported with readr (v2.1.4) and processed using dplyr (v1.1.4) and tidyr (v1.3.0), with removal of contaminants and low-confidence entries. Phosphosite-level specificity was preserved by generating unique feature identifiers combining gene symbol and phosphorylation site annotation (*Gene|site_label*), and numeric columns were explicitly coerced to the numeric format. Differential abundance or phosphorylation was assessed for each treatment condition (BX795, Dab, and their combination) relative to the control, yielding log_2_ fold changes (log_2_FCs) and associated p-values, with multiple testing correction applied using the Benjamini–Hochberg method. Volcano plots (log_2_FC versus −log_10_(p-value)) and Venn diagrams of significantly regulated features were generated using ggplot2 (v3.4.4), ggrepel (v0.9.4), and ggVennDiagram (v1.2.2). Gene set enrichment analysis (GSEA) was performed with fgsea (v1.28.0) using a signed ranking metric (sign(log_2_FC) × −log_10_(p-value)) and curated Kyoto Encyclopedia of Genes and Genomes (KEGG) and MSigDB Hallmark gene sets, reporting normalized enrichment scores (NESs) with FDR-adjusted q-values. In addition, enrichment heatmaps were produced with pheatmap (v1.0.12). All figures were exported at publication-quality resolution.

### Cell cycle analysis

Mutant *BRAF* V600E ATC cells (8505C and SW1736) were treated overnight with DMSO (control), BX795 (2.5 µM), Dab (2.5 µM), or their combination (BX795 + Dab). Then, cells were trypsinized and fixed using pre-chilled 70% ethanol, followed by overnight incubation at –80 °C. The next day, cells were thawed and centrifuged at 1500 rpm for 5 min, washed with PBS, and stained with FxCycle^TM^ PI/RNase staining solution (Thermo Fisher Scientific) for 30 min at room temperature. After incubation, the cell cycle and DNA content (2N or 4N) were analyzed with a BD Accuri C6 Plus flow cytometer (BD Biosciences, San Jose, CA, USA). Experiments were performed in triplicate, and the results are reported compared to the control.

### Apoptosis assay

Apoptotic cells were identified using the Annexin V/Dead Cell Apoptosis Kit (BD Biosciences) as per the manufacturer’s protocol. Briefly, mutated *BRAF* V600E ATC cells (8505C and SW1736) were treated for 48 h with DMSO (control), BX795 (2.5 µM), Dab (2.5 µM), or their combination (BX795 + Dab). Then, cells were harvested, trypsinized, and washed with ice-cold PBS. The single-cell suspension was resuspended in 100 μL of 1× annexin-binding buffer with 5 µL each of Annexin V and propidium iodide (PI), followed by incubation for 20 min at room temperature. Annexin V/PI-stained cells were measured using a BD Accuri C6 Plus flow cytometer (BD Biosciences). The relative fold change was determined compared to the control.

### Reactive oxygen species (ROS) measurement

Cellular ROS were measured using DCHFDA staining. Mutant *BRAF* V600E ATC cells (8505C and SW1736) were treated 24h with DMSO (control), BX795 (2.5 µM), Dab (2.5 µM), or their combination (BX795 + Dab). After treatment, cells were trypsinized, washed with PBS, and stained with DCHFDA (5 µM) for 30 min in the dark. After incubation, DCHFD fluorescence was measured using a BD Accuri C6 Plus flow cytometer (BD Biosciences). The results are presented as relative fold change compared to the control.

Mitochondrial ROS production was measured using Mito-SOX™ Red mitochondrial superoxide indicator. After treatment overnight, cells were harvested, trypsinized, washed with PBS, and stained with 2.5 μM Mito-SOX Red for 20 min at 37 °C. Mito-SOX^TM^ fluorescence was measured using a BD Accuri C6 Plus flow cytometer (BD Biosciences). The results are presented as relative fold change compared to the control.

### Mitochondrial content measurement

Mitochondria were visualized using MitoTracker Green FM (Thermo Fisher Scientific), which labels mitochondria independently of the membrane potential. Mutant *BRAF* V600E ATC cells (8505C and SW1736) were seeded onto sterile coverslips in 12-well plates; allowed to adhere overnight; and treated with DMSO (control), BX795 (2.5 µM), Dab (2.5 µM), or their combination (BX795 + Dab). After treatment, cells were stained with 50 nM MitoTracker Green for 30 min at 37 °C in the dark. Then, cells were washed with PBS, and live cell fluorescence imaging was acquired using a SP8 confocal microscope (Leica, Wetzlar, Germany) at 400× magnification under identical acquisition settings for all samples at multiple sites on each slide. Fluorescence intensity was quantified using ImageJ.

In addition, single-cell suspensions of treated cells were stained in triplicate with 50 nM MitoTracker Green FM for 30 min at 37 °C in the dark. Cellular fluorescence was measured using a BD Accuri C6 Plus flow cytometer (BD Biosciences). The results are presented as the relative fold change compared to the control.

### Measurement of mitochondrial transmembrane potential

The mitochondrial membrane potential was analyzed by using the Image-iT™ TMRM Reagent (Thermo Fisher Scientific). Mutant *BRAF* V600E ATC cells (8505C and SW1736) were treated overnight in triplicate with DMSO (control), BX795 (2.5 µM), Dab (2.5 µM), or their combination (BX795 + Dab). After treatment, cells were trypsinized, and single-cell suspensions were stained with TMRM (100 nM) for 30 min at 37 °C in the dark. Following treatment, cells were washed and resuspended in ice-cold PBS. Fluorescence was measured using a BD Accuri C6 Plus flow cytometer (BD Biosciences). The results are presented as the relative fold change compared to the control.

Mitochondrial depolarization was measured with the potentiometric dye JC-1. After treatment 24h with DMSO (control), BX795 (2.5 µM), Dab (2.5 µM), or their combination (BX795 + Dab), mutant *BRAF* V600E ATC cells (8505C and SW1736) were stained with JC-1 (2 μM) for 30 min at 37 °C in the dark, washed, and resuspended in ice-cold PBS. Red and green fluorescence was measured using a BD Accuri C6 plus flow cytometer (BD Biosciences). The results are presented as the relative ratio of red versus green fluorescence compared to the control.

### Measurement of the oxygen consumption rate (OCR)

The OCR was measured using a Seahorse XFe24 Extracellular Flux Analyzer (Seahorse Bioscience, Santa Clara, CA). Mutant *BRAF* V600E ATC cells (8505C and SW1736) were seeded evenly in XF24 microplates; allowed to adhere for 24 h; and then treated for an additional 24 h with DMSO (control), BX795 (2.5 µM), Dab (2.5 µM), or their combination (BX795 + Dab). Measurements were performed in XF assay medium containing 1 mM sodium pyruvate, 2 mM glutamine, and 10 mM glucose under basal conditions. After washing with assay medium, cells were incubated at 37 °C in a non-CO₂ incubator for 30 min. The baseline OCR was recorded, followed by sequential injections of oligomycin (1 µM, an ATP synthase inhibitor), FCCP (4 µM, a mitochondrial uncoupler), rotenone (1 µM, a complex I inhibitor), and antimycin A (1 µM, a complex III inhibitor) using Seahorse XFe24 Extracellular Flux Analyzer (Seahorse Bioscience). Mitochondrial function parameters were calculated as follows: basal OCR = baseline OCR − OCR after rotenone/antimycin A; ATP-linked OCR = basal OCR − OCR after oligomycin; maximal respiration = OCR after FCCP − OCR after rotenone/antimycin A. The assay involved 3 min of mixing, 3 min of waiting, and 3 min of measurement. The data were normalized to the total protein content and are presented as the mean ± standard deviation (SD).

### Western blot analysis

Lysates from mutant *BRAF* V600E ATC cells (8505C and SW1736) were prepared in radioimmunoprecipitation assay (RIPA) buffer supplemented with phosphatase and protease inhibitors. Protein concentrations were determined using the Pierce™ BCA Protein Assay Kit (Thermo Fisher Scientific). Equal amounts of protein were separated by electrophoresis on 4%–15% precast sodium dodecyl sulfate–polyacrylamide gel electrophoresis (SDS-PAGE) gels and transferred to nitrocellulose membranes. Membranes were incubated overnight at 4 °C with primary antibodies (Table S1), followed by horseradish peroxidase (HRP)-conjugated secondary antibodies (Table S1). Protein bands were visualized using enhanced chemiluminescence (ECL) and imaged with a Bio-Rad ChemiDoc™ MP Imaging System (Bio-Rad, Hercules, CA, USA). Band intensities were quantified using ImageJ.

### Immunofluorescence

Mutant *BRAF* V600E ATC cells (8505C and SW1736) were treated overnight with DMSO (control), BX795 (2.5 µM), Dab (2.5 µM), or their combination (BX795 + Dab). Then, cells were washed with PBS, fixed 3 × 15 min using chilled methanol, washed 3 × 5 min with PBS, permeabilized by PBS containing 0.2%Triton X-100, and finally washed three times with PBS. Cells were incubated with 1% bovine serum albumin containing 0.02% Tween 20 for 30 min at room temperature (to block nonspecific protein binding), followed by incubation overnight with the pH_2_AX primary antibody (Table S1) at 4 °C. The next day, cells were washed three times with PBS containing 0.02% Tween 20 (PBST) and incubated with fluorescently labelled secondary antibodies (Table S1) for 2 h at room temperature. Finally, cells were washed with PBST and counterstained with mounting media containing DAPI. Labelled proteins were observed using an SP8 confocal microscope (Leica) at 400× magnification. The number of H2AX foci among 25 cells was counted for each treatment. The results are presented as the number of foci (mean ± standard error of the mean [SEM]).

### Orthotopic ATC xenograft model

For the orthotopic ATC model, 1 × 10^6^ 8505C-Luc2 cells (Luc2 denotes cells with stable expression of a luciferase reporter) were implanted into the thyroid gland of male and female *NOD Cg-Prkdcs^cid^ Il2rg^tm1WjI^/SzJ* mice. All animal procedures were approved by the Stanford Administrative Panel on Laboratory Animal Care (APLAC; Protocol #34767) and conducted in accordance with institutional and federal guidelines. Tumor luminescence was measured using the Lago (Spectral Instruments Imaging, Tucson, AZ, USA) after intraperitoneal luciferin injection (150 mg/kg). One week after orthotopic implantation, 10 mice (5 males and 5 females) were randomized into four treatment groups: vehicle control (CM-cellulose), BX795 (100 mg/kg), Dab (15 mg/kg), and BX795 (100 mg/kg) + Dab (15 mg/kg). BX795 and Dab were solubilized in methyl cellulose (0.5% in double-distilled water) with sonication (Picoruptor, Diagenode Inc, Devville, NJ, USA) at 4 °C. They were administered orally 5 days/week for 2 weeks with continuous monitoring for toxicity. Mice were imaged and weighed weekly. After treatment, all mice were euthanized using CO_2_ inhalation. Tumor, lung, and liver samples were collected after euthanasia. Tumor size and weight were recorded, and thyroid tumor, lung, liver, and kidney samples were collected for histopathological evaluation.

### Statistical analyses

The data are presented as the mean ± SD, with the details for each analysis provided in the figure legends. Note that the *in vivo* results and number of H_2_AX foci are presented as the mean ± SEM. All *in vitro* experiments were performed in at least three independent experiments. Analysis of variance (ANOVA) was used to determine the differences between groups. GraphPad Prism version 10 (GraphPad Software, San Diego, CA) was used for all statistical analyses. The level of statistical significance is indicated as follows: ns is nonsignificant; *p < 0.05; **p < 0.01; ***p < 0.001; and ****p < 0.0001.

## RESULTS

### Functional dependency on PDPK1 in ATC

To assess PDPK1 as a therapeutic vulnerability in ATC, we integrated DepMap CRISPR Chronos dependency scores with PDPK1 expression across cancer lineages. PDPK1 showed broad essentiality (median Chronos ≈ –0.88), with thyroid cancers clustering among the highly dependent groups, comparable to bowel and pancreatic cancers. Within the thyroid lineage, PDPK1 transcript levels were similar across medullary, well-differentiated, and anaplastic thyroid cancer cell lines (Fig. S1A–B). In contrast, functional dependency analysis revealed marked PDPK1 dependence in both ATC (Chronos –0.65 to –1.15; median –0.84) and WDTC (–0.606 to –1.15; median –0.98) models (Fig. S1C). Dependency aligned with oncogenic drivers: the *BRAF* V600E-mutant BHT101 line was most sensitive (Chronos –1.15), followed by other *BRAF* V600E-mutant lines (BCPAP, 8305C, 8505C), all showing high PDPK1 expression and strong dependency (Table S4). PDPK1 expression correlated tightly with Chronos dependency (r = 0.97, p < 0.03), reinforcing its functional relevance in thyroid cancer.

PDPK1 activity is regulated predominantly at the post-translational level; therefore, messenger RNA (mRNA) or total protein abundance may not reliably reflect its functional status (46). The critical activation step for PDPK1 is autophosphorylation at Ser241, which generates the catalytically competent form required for downstream phosphorylation of AKT (Thr308), SGK, and other AGC kinases. Because only the phosphorylated, active pool (p-PDPK1 S^241^) propagates PI3K/AKT signaling, quantifying this phosphorylated site provides a direct measure of pathway engagement and oncogenic dependency. Therefore, we first characterized total and phospho-PDPK1 in a panel of thyroid cancer cell lines. PDPK1 expression and phosphorylation varied in the cell lines with known mutations in genes involved in the MAPK and PI3K/AKT pathways. Cell lines harboring *PTEN* loss (FTC236, FTC133, and C643) and *BRAF* V600E mutations (SW1736, 8505C, and BCPAP), as well as wild-type *BRAF* cell lines (THJ29T and C643) and a *PIK3CA*-mutant/*MKRN1–BRAF* fusion cell line (THJ16T), all exhibited measurable PDPK1 expression (Fig. 1A). The phospho-PDPK1/PDPK1 ratio was markedly elevated in the FTC236, C643, THJ16T, BCPAP, and 8505C cell lines relative to the HEK293 cell line (control), with the THJ16T cell line demonstrating the highest PDPK1 phosphorylation, followed by the homozygous *BRAF* V600E–mutant 8505C and BCPAP cell lines (Fig. 1A).

**Fig. 1:**
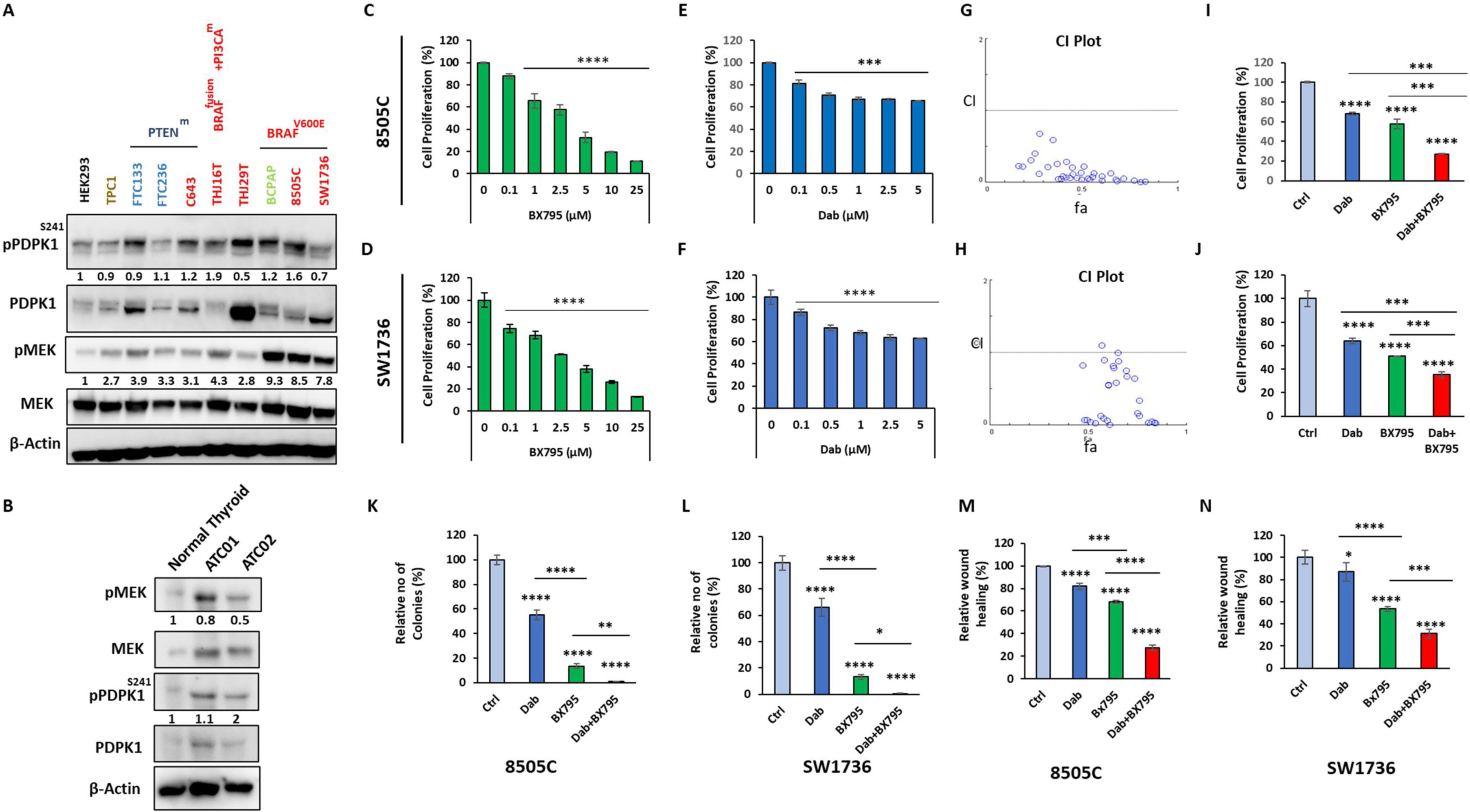
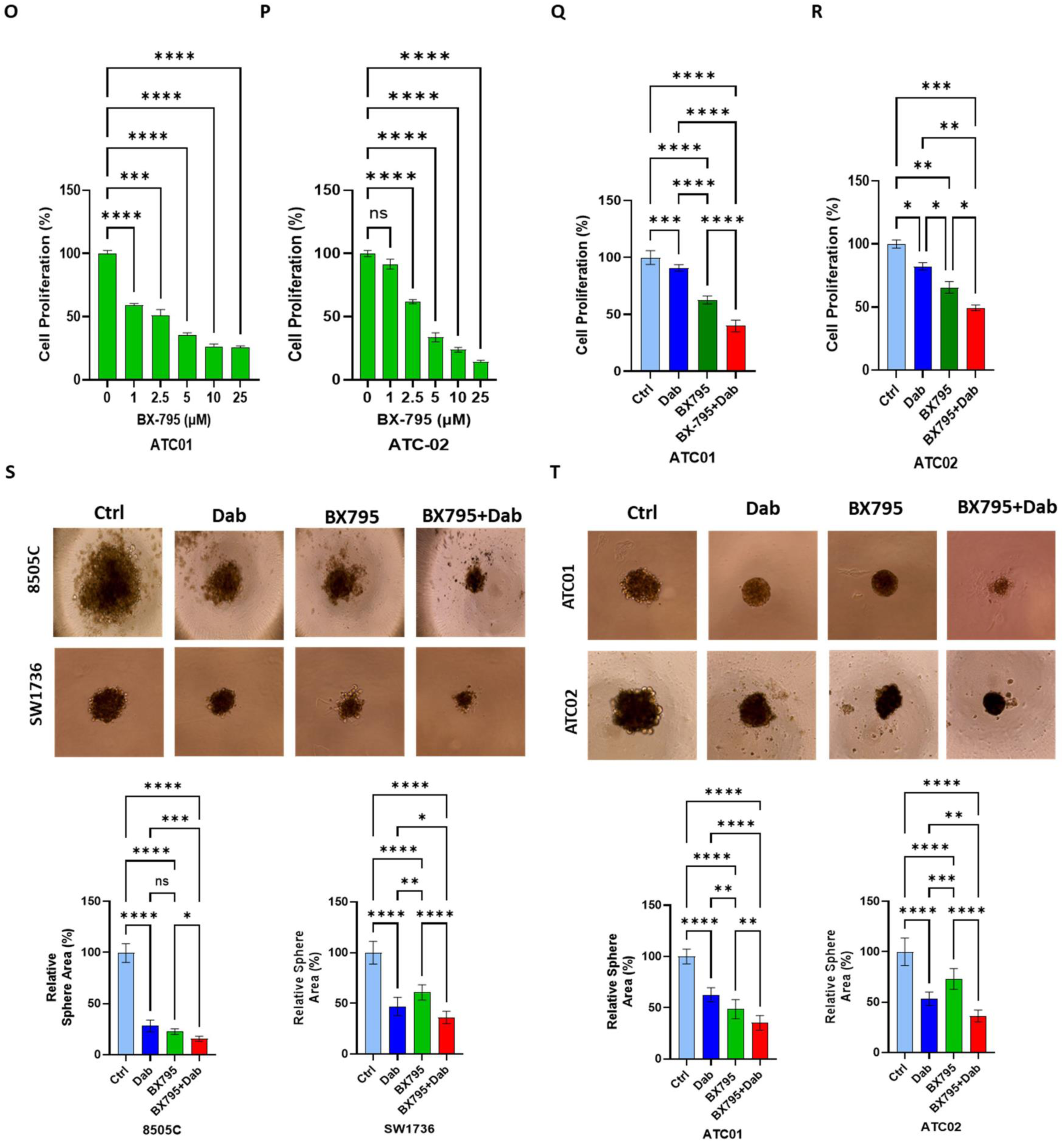
Combined PDPK1 and BRAF V600E inhibition has synergistic anticancer activity *in vitro* and *ex vivo*. (A) Immunobloting of pPDPK1, PDPK1, pMEK, and MEK in thyroid cancer lines: papillary thyroid cancer (PTC, brown color) (TPC1), follicular thyroid cancer (FTC, blue color) (FTC-133 and FTC-236), poorly differentiated thyroid cancer (PDTC, green color) (BCPAP), *BRAF* wild type anaplastic thyroid cancer (ATC) (red color) (C643 and THJ29T), *BRAF*V600E–mutant ATC (8505C, SW1736, and THJ16T), and HEK293 cells using β-actin as a loading control. The relative ratio of pPDPK1 to total PDPK1 and pMEK to total MEK is listed below each phospho-protein compared to HEK293 cells. (B) Immunoblotting of phosphoPDPK1, total PDPK1 in untreated and dabrafenib (Dab) and trametinib–treated patient derived ATC and normal thyroid tissues using β-actin as a loading control. The relative ratio of pPDPK1 to total PDPK1 and pMEK to total MEK is listed below each phospho-protein compared to normal thyroid tissue. (C–D) Dose-dependent inhibition of cell proliferation with BX795 treatment in 8505C and SW1736 cells after 48 h. (E–F) Dose-dependent inhibition of cell proliferation with Dab treatment in 8505C and SW1736 ATC cells after 48 h. (G–H) Analysis of the combination index (CI) using the CompuSyn software for BX795, Dab, and their combination in *BRAF* V600E–mutant ATC cell lines. CI < 1 indicates synergism, and fa denotes fraction affected. (I–J) The effect of BX795 (2.5 µM), Dab (2.5 µM), and their combination on cellular proliferation in 8505C and SW1736 cells after 48 h. (K–L) Colony-formation assay in 8505C and SW1736 cells treated with BX795 (2.5 µM), Dab (2.5 µM), or their combination. Quantification was performed using ImageJ. (M–N) Cellular migration as measured with a wound-healing assay in 8505C and SW1736 cells treated with BX795 (2.5 µM), Dab (2.5 µM), or their combination. Quantification was performed using the ImageJ software. (O–P) Concentration-dependent effects of BX795 treatment on proliferation of patient-derived ATC cells (ATC01 and ATC02). ATC01 was derived from a treatment-naïve patient tumor positive for *BRAF* V600E mutation, and ATC02 was derived from a residual tumor in a patient with an exceptional treatment response to Dab and trametinib, and the tumor was positive for *BRAF* V600E mutation. (Q–R) The effects of treatment with BX795 (2.5 µM), Dab (2.5 µM), or their combination on the proliferation of patient-derived ATC cells (ATC01 and ATC02). (S) The effect of treatment with BX795 (2.5 µM), Dab (2.5 µM), or their combination on ATC spheroids in *BRAF* V600E–mutant in the 8505C and SW1736 cell lines. (T) The effect of treatment with BX795 (2.5 µM), Dab (2.5 µM), or their combination on patient-derived ATC spheroids (ATC01 and ATC02). All data are presented as the mean ± standard deviation. Statistical significance is indicated as ns = nonsignificant; *p < 0.05; **p < 0.01; ***p < 0.001; ****p < 0.0001.

Consistent with MAPK pathway activation, all thyroid cancer cell lines evaluated showed high MEK phosphorylation, with the *BRAF* V600E–mutant cell lines (BCPAP, 8505C, and SW1736) exhibiting the highest phospho-MEK levels (Fig. 1A). The phospho-MEK/MEK ratio was lower in the two patient-derived *BRAF* V600E–mutant ATC cell lines (ATC01 and ATC02) compared to normal thyroid tissue, but total MEK levels were much higher in the patient-derived ATC cell lines (Fig. 1B). The phospho-PDPK1/PDPK1 ratio was higher in the ATC02 cell line, which was derived from a residual tumor resected after an exceptional response to combined Dab and trametinib treatment, compared to the ATC01 cell line (derived tumor from a treatment-naïve patient) or normal thyroid tissue. We used the *BRAF* V600E–mutant ATC cell lines (8505C and SW1736) with relatively high and low phospho-PDPK1 levels and the patient-derived cell lines (ATC01 and ATC02) to test the therapeutic efficacy of targeting PDPK1.

### Combined PDPK1 and BRAF V600E inhibition has synergistic anticancer activity *in vitro* and *ex vivo*

We evaluated drug response data from our previous drug screening studies for PDPK1 inhibitors in three ATC cell lines (8505C, THJ-11T, and THJ-16T), where BX795 emerged as the most consistent and broadly active compound based on a half-maximal activity concentration (AC_50_) < 10 µM and **a** curve class up to –2.2. BX795 was the only agent to meet the criteria in all three ATC cell lines, whereas BX-912, OSU-03012, and GSK-2334470 met the criteria in two ATC cell lines, and DTXSID90678515 met the criteria in one ATC cell line (Fig. S1D). Therefore, we used BX795 to inhibit PDPK1 phosphorylation. In addition, Integrated DepMap PRISM analyses indicated that thyroid cancer cells are sensitive to BX795 treatment (Fig. S1E), further supporting that the PDPK1 gene is essential. BX795 treatment significantly inhibited cell proliferation in *BRAF* V600E–mutant ATC cell lines in a dose-dependent manner (Fig. 1C). BX795 treatment reduced phosphorylation of PDPK1 at Ser241 and PI3KAKT/mTOR signaling, with reduced S6K levels and upregulation of H_2_AX phosphorylation (Fig. S1F). To further validate the essentiality of PDPK1 and specificity of BX795 treatment, we knocked down PDPK1 expression. PDPK1 knockdown reduced total and phospho-PDPK1, changes associated with inhibition of cellular proliferation (Fig. S1G). When combined with BX795 treatment to inhibit PDPK1 phosphorylation at Ser241, there was further inhibition of cell proliferation compared to the control (Fig. S1G). Collectively, the DepMap datasets and *in vitro* results suggest that targeting PDPK1 for ATC therapy is a promising approach, especially in *BRAF* V600E–mutant ATC, where resistance to combined BRAF and MEK inhibition is common.

PDPK1 inhibition with BX795 and BRAF V600E inhibition with Dab alone inhibited ATC cellular proliferation (Fig. 1C–F). Combination BX795 and Dab treatment showed a synergistic antiproliferative effect in *BRAF* V600E–mutant ATC cell lines at multiple dose combinations (Fig. 1G–J) based on a CI < 1. We selected combination concentrations (2.5 µM BX795 and 2.5 µM Dab) that were synergistic and clinically achievable for the follow-up studies. Combination BX795 and Dab treatment significantly inhibited the size and number of ATC colonies and cellular migration (Figs. 1K–N and S1H–I) compared to either agent alone or the vehicle control. BX795 alone and combination BX795 and Dab treatment in patient-derived *BRAF* V600E–mutant ATC cells also significantly inhibited cellular proliferation (Fig. 1O–R).

We next tested the anticancer activity of combination BX795 and Dab treatment in *ex vivo* (3D culture) model of *BRAF* V600E–mutant ATC cell lines (8505C and SW1736) and two patient-derived *BRAF* V600E–mutant ATC cells (ATC01 and ATC02) that better recapitulate the molecular features of human ATC and response to treatment. Dab treatment alone reduced the spheroid size by ∼72% (8505C) and ∼56% (SW1736), and BX795 treatment alone reduced the spheroid size by ∼78% (8505C) and ∼40% (SW1736) compared to the vehicle control; combination treatment reduced the spheroid size significantly more than either agent alone (Fig. 1S–T). Similarly, in patient-derived ATC01 and ATC02 spheroids, combination treatment reduced the spheroid size significantly more than either agent alone or the vehicle control (Fig. 1S–T).

### Combined PDPK1 and BRAF V600E inhibition has anticancer activity and is safe in *in vivo* orthotopic model of ATC

We next tested combination BX795 and Dab treatment in an orthotopic mouse model using the ATC *BRAF* V600E–mutant 8505C ATC cell line. BX795 and Dab treatment alone reduced the tumor volume by 18% and 39%, respectively, compared to the vehicle control, whereas combination BX795 and Dab treatment reduced the tumor volume by 55% compared to the vehicle control (Figs. 2A and S2A–B). Combination BX795 and Dab treatment also significantly reduced the tumor weight compared to the vehicle control in both genders (Fig. 2B; S2C-D). The total body tumor luminescence measurement was significantly lower after treatment for 2 weeks, and the growth rate was lower for the combination BX795 and Dab treatment group compared to the vehicle control. Moreover, there were no metastases in any of the treatment groups (Fig. 2C–D). There was no statistically significant difference in mouse weight among treatment groups, and there were no treatment-related toxicities (Fig. 2E).

**Fig. 2:**
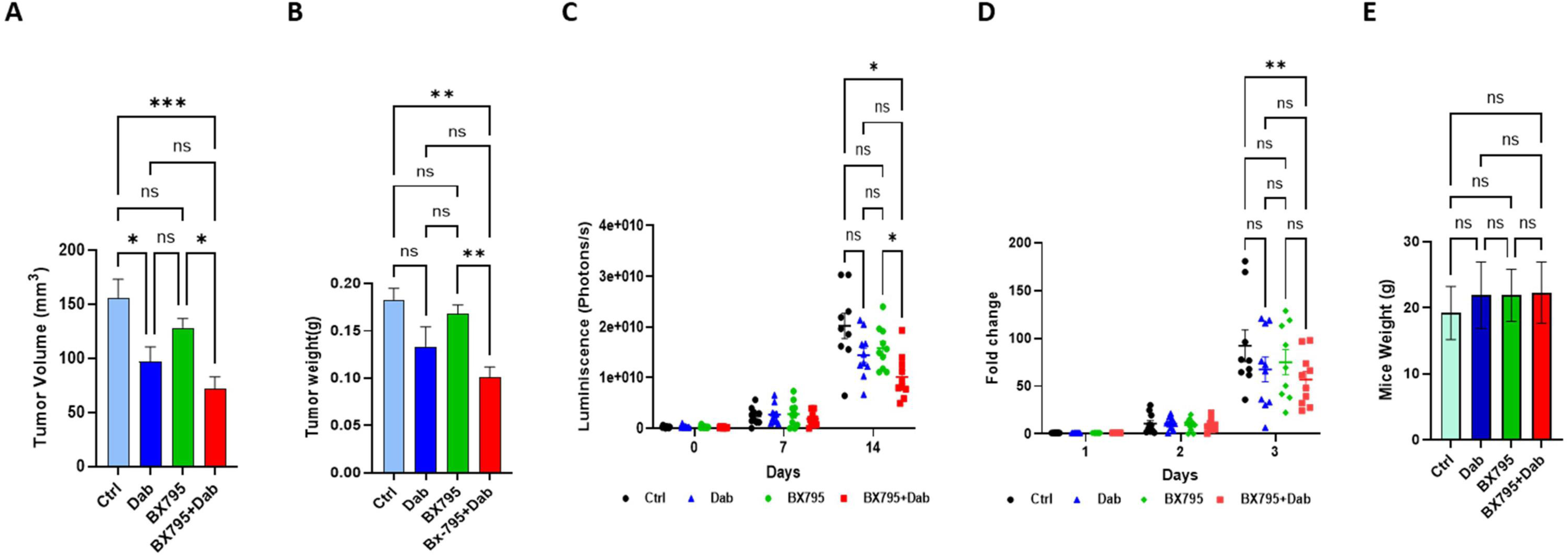
Effect of combination BX795 and dabrafenib (Dab) treatment in an orthotopic ATC *in vivo* model. Mice were treated with BX795 (100 mg/kg), Dab (15 mg/kg), or their combination 5 days per week. After 2 weeks, tumors were excised and their volume (mm^3^) (A) and weight (B) were measured. (C–D) *In vivo* tumor growth was analyzed by tumor luminescence analysis (photons/s) (C), and the fold change of each tumor was determined relative to the luminescence intensity of the tumor site at day 0 (D). Comparisons were made across treatment groups at days 0, 7, and 14. (E) Body weight of mice on day 14 of the study. All data are presented as mean ± standard error of the mean. Statistical significance is indicated as ns = nonsignificant; *p < 0.05; **p < 0.01; ***p < 0.001.

### Combined PDPK1 and BRAF V600E inhibition induces apoptosis

To investigate the mechanism of the synergistic anticancer activity of combined PDPK1 and BRAF V600E inhibition, we performed TMT-based, label-intensity proteomic analysis with BX795, Dab, or combination treatment. The results showed distinct pathway-selective effects with monotherapy and a synergistic signaling convergence with combination treatment. Volcano plot analysis demonstrated that BX795 treatment preferentially suppressed proteins involved in PDPK1/AKT/mTOR signaling, ribosomal biogenesis, mitochondrial regulators, and cytoskeletal organization, while inducing checkpoint- and stress-associated regulators (Fig. 3A). In contrast, Dab treatment predominantly repressed MAPK-dependent proliferative and mitotic programs, with compensatory activation of nuclear structural and transcriptional related proteins (Fig. 3B). Combination treatment produced the greatest magnitude and breadth of proteomic remodeling, marked by strong suppression of epithelial–mesenchymal transition (EMT)/metastasis-associated proteins, translational machinery, and chromatin regulators, together with induction of DNA damage, mitotic spindle destabilization, and apoptotic execution factors, consistent with mitotic catastrophe (Fig. 3C). Venn diagram analysis revealed a conserved core of 134 shared total proteins and 104 shared phosphoproteins across all three treatment groups, defining a common stress-adaptive signaling module, while combination treatment uniquely altered 108 total proteins and 40 phosphoproteins, indicating novel vulnerabilities unmasked only under dual pathway inhibition (Fig. 3D). At the signaling level, phosphoproteomic analysis confirmed that BX795 suppressed PI3K/AKT/mTOR-dependent phosphorylation, Dab inhibited MEK/ERK and mitotic phospho-networks, and combination treatment resulted in simultaneous shutdown of both pathways, eliminating compensatory feedback upregulation. GSEA further demonstrated that BX795 negatively enriched the PI3K/AKT/mTOR, mitotic spindle, oxidative phosphorylation, WNT/β-catenin, and DNA repair pathways, while activating G2/M checkpoint inflammatory stress and MAPK signaling. Dab suppressed MAPK-driven mitotic and metabolic programs with activation of DNA repair, G2M checkpoints, EMT, p53, and oxidative phosphorylation checkpoint pathways (Fig. 3E–F). Strikingly, combination treatment produced the most profound pathway-level effects, with coordinated suppression of PI3K/AKT/mTOR, MAPK, WNT, mitosis, oxidative phosphorylation, and DNA repair, alongside enrichment of p53, G2/M checkpoint, and inflammatory response pathways, and repression of EMT signaling (Fig. 3G). Collectively, these data establish that dual PDPK1 and BRAF V600E inhibition drives a synergistic, multi-layered inhibition of survival, translational, cytoskeletal, mitotic, metabolic, and DNA repair networks, leading to cell death.

**Fig. 3:**
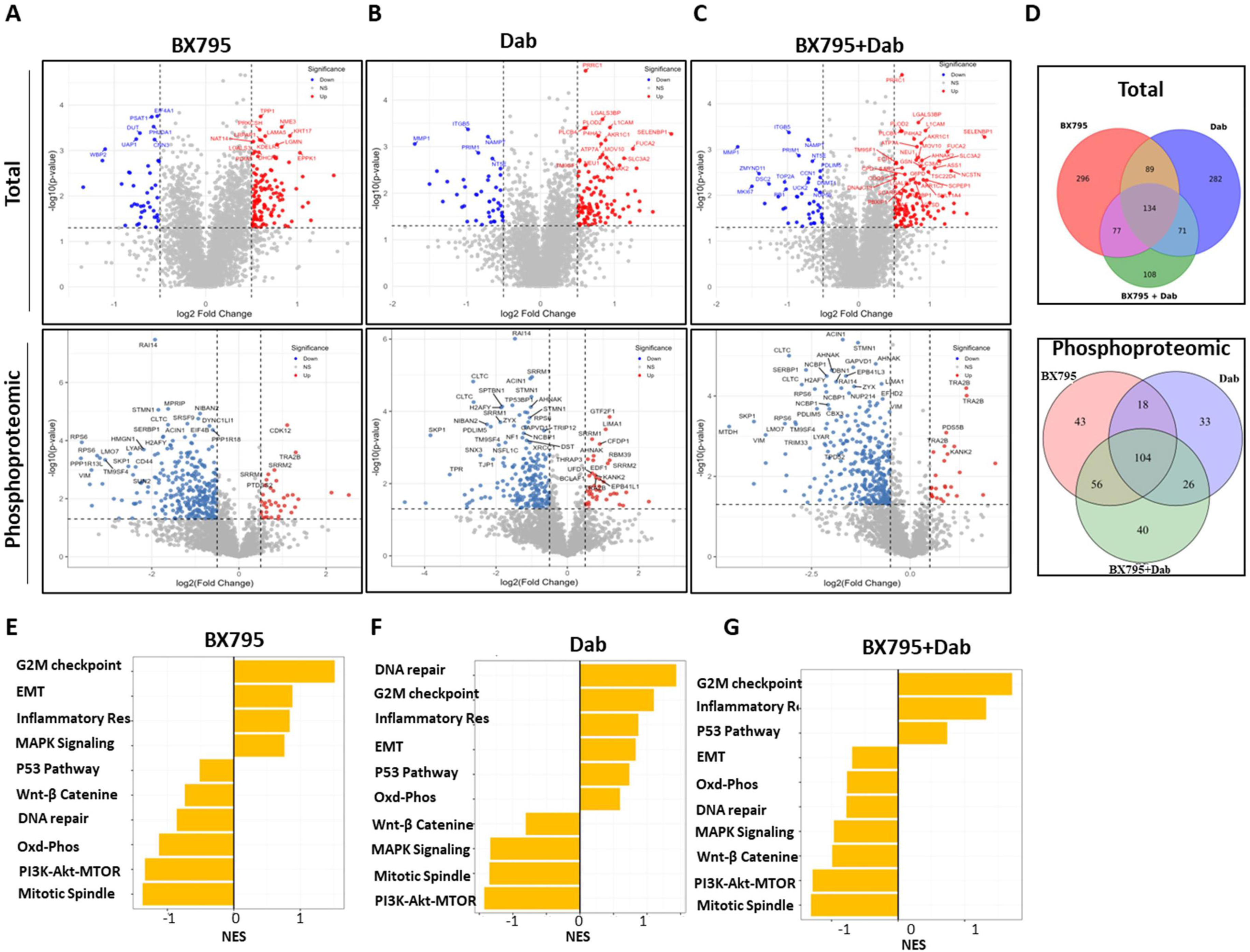
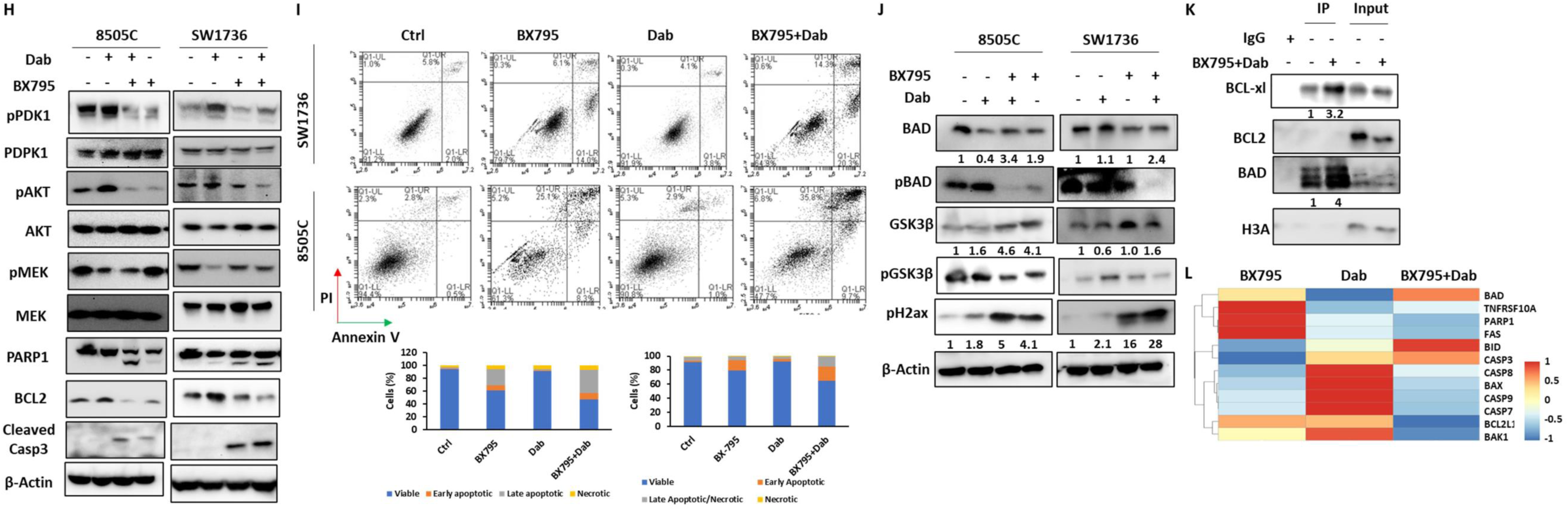
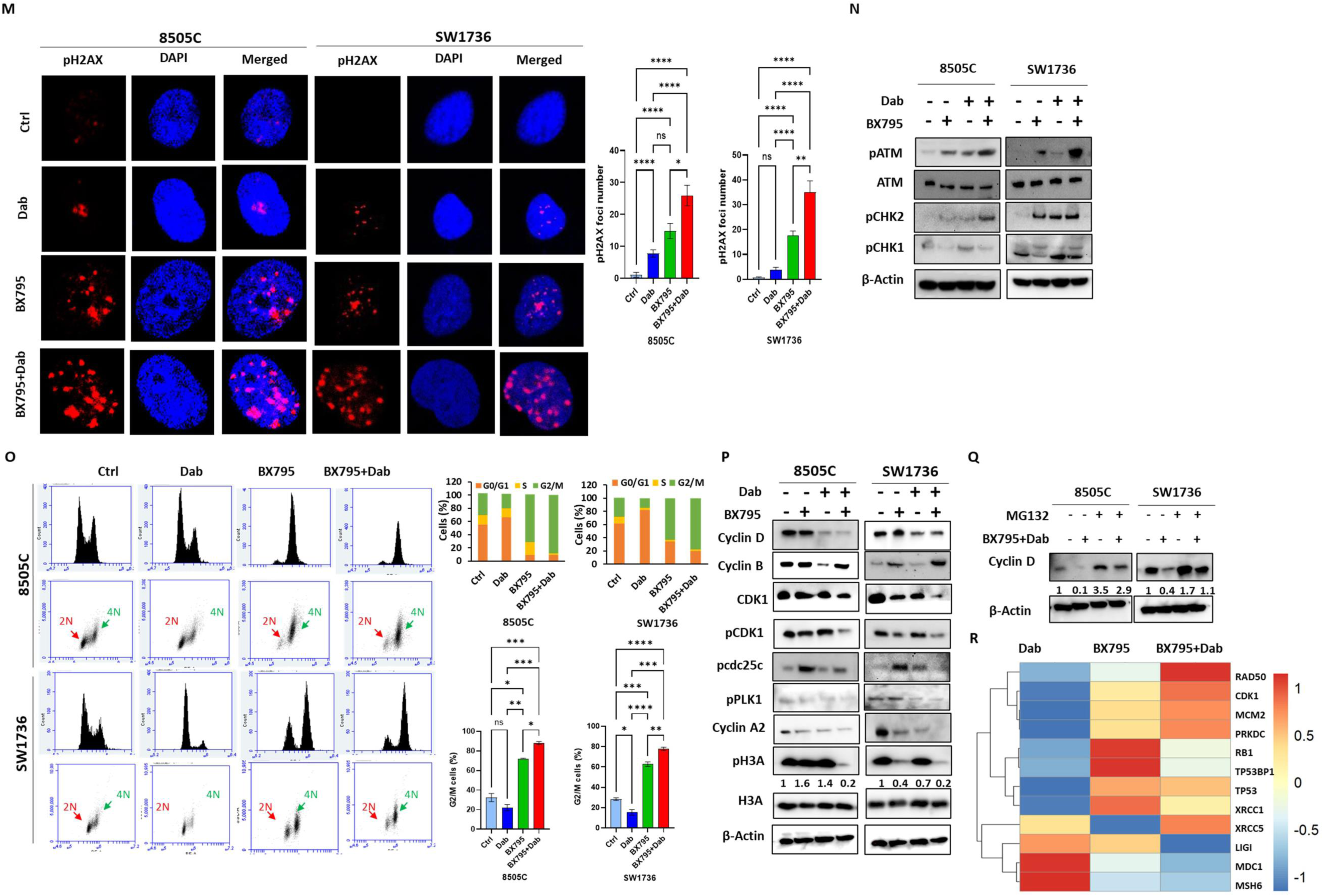
Combined PDPK1 and BRAF V600E inhibition induces apoptosis and DNA damage in *BRAF* V600E–mutant anaplastic thyroid cancer (ATC). (A) Mutated *BRAF* V600E ATC cells (8505C) were treated for 48 h with BX795 (2.5 µM), dabrafenib (Dab) (2.5 µM), or their combination. Then, total and phosphorylated protein levels were measured using mass spectrometry. (A–C) The volcano plots show significantly upregulated proteins in red, significantly downregulated proteins in blue (p < 0.05), and non-significant proteins in gray. The x-axis represents log_2_ fold change, and the y-axis shows –log_10_(p-value). The dashed lines indicate thresholds for statistical significance and fold change. (D) The Venn diagrams shows shared and treatment-specific differentially expressed proteins and phosphorylated proteins across the treatment groups (p < 0.05). The overlapping regions represent a conserved core proteomic response, while the non-overlapping regions indicate pathway-specific effects of PDPK1 or BRAF inhibition. (E–G) Gene set enrichment analysis (GSEA) of ranked phosphorylated site changes following BX795 (E), Dab (F), or combination (G) treatment. The bar plots show normalized enrichment scores (NES) for Hallmark Kyoto Encyclopedia of Genes and Genomes (KEGG) signaling pathways. (H) Western blot analysis of the effect of BX795 (2.5 µM), Dab (2.5 µM), or combination treatment on *BRAF* V600E–mutant ATC cell lines (8505C and SW1736) after 48 h. pMEK, total MEK, pAKT^308^, total AKT, pPDPK1^S241^, PARP-1, BCL2, and caspase-3 protein levels are shown, with β-actin used as a loading control. (I) Flow cytometry results of Annexin V/PI staining with BX795 (2.5 µM), Dab (2.5 µM), or combination treatment. (J) Western blot analysis of apoptosis regulatory proteins 40 h after treatment. pBAD^S112^, BAD, BCL2, pGSK3β^Ser9^, GSK3β, and the double-stranded DNA damage marker pH2AX^S139^ are shown, with β-actin used as a loading control. Protein band density was measured and normalized to β-actin. The phosphorylated to total protein ratios were measured, and the values are listed above the phosphoprotein band. Protein band density of BCL-xL and pH2AX^S139^ were also measured and normalized to β-actin. (K) Immunoprecipitation of BCL-xL and BCL2 with BAD 30 h after treatment with BX795 and Dab in *BRAF* V600E–mutant ATC cell lines. Protein band density was measured and normalized to the input protein band. The ratio of immunoprecipitated to total input protein was measured and is listed below the protein blot. P = phosphorylation. The superscript indicates the amino acid phosphorylation site. (L) The heatmap shows z-scored log_2_-transformed abundance of apoptosis-related proteins for the BX795, Dab, and combination treatment groups in 8505C ATC cells. The proteins are displayed in sequential order—BAD, TNFRSF10A, PARP1, FAS, BID, CASP3, CASP8, BAX, CASP9, CASP7, BCL2L1, and BAK1—with hierarchical clustering applied to the rows. (M) The effect of BX795 (2.5 µM), Dab (2.5 µM), or combination treatment for 16 h on γH2AX foci formation in the 8505C and SW1736 ATC cell lines using immunofluorescence. γH2AX was labelled with Alexa Fluor™ 546 secondary antibody, with DAPI used for nuclear staining. A Zeiss LSM 800 confocal microscope was used to examine the cells (400× magnification). The number of foci per cell was quantified using ImageJ. All data are presented as the mean ± standard error of the mean of 25 cells. Statistical significance is indicated as ns = nonsignificant; *p < 0.05; **p < 0.01; ***p<0.001; ****p < 0.0001. (N) Western blot analysis of the effect of BX795 (2.5 µM), Dab (2.5 µM), or combination treatment for 16 h on DNA damage–dependent proteins (pATM, total ATM, pCHK2, Total CHK2, and pCHK1). β-Actin was used as a loading control. (O) Cell cycle analysis after BX795 (2.5 µM), Dab (2.5 µM), or combination treatment for 16 h in *BRAF* V600E–mutant ATC cell lines (8505C and SW1736). Cells were stained with propidium iodide (PI) and analyzed by fluorescence-activated cell sorting. The DNA content was measured, 2N (diploid) and 4N (tetraploid), based on the PI-stained DNA content. (P) The effect of BX795 (2.5 µM), Dab (2.5 µM), or combination treatment for 16 h on DNA repair proteins—cyclin D, cyclin B, pcdc25c, pCDK1, total CDK1, pH3A, H3A, and cyclin A2—by immunoblotting. β-actin was used as a loading control. The ratio of phospho-H3A to total H3A was measured and is listed below the phospho-H3A protein band. All data are presented as the mean ± standard deviation. Statistical significance is indicated as ns = nonsignificant; *p < 0.05; **p < 0.01; ***p < 0.001; ****p < 0.0001. (Q) Analysis of the effect of combination BX795 (2.5 µM) and Dab (2.5 µM) treatment for 16 h followed by proteasomal degradation inhibition for 6 h (MG132, 10 µM). Protein band density was measured and normalized to β-actin. (R) The heatmap shows z-scored log2-transformed phosphorylation levels of DNA damage response, DNA repair, and cell-cycle regulatory proteins for the BX795, Dab, and combination treatment groups in 8505C ATC cell line. Proteins are displayed in sequential order as RAD50, CDK1, MCM2, PRKDC, RB1, TP53BP1, TP53, XRCC1, XRCC5, LIG1, MDC1, and MSH6, with supervised hierarchical clustering applied to the rows. The color scale represents relative protein levels (red = higher; blue = lower).

To validate the proteomic results, we analyzed the canonical signaling pathways targeted by BX795 and Dab, and cell death pathways. As expected, Dab reduced MEK phosphorylation, and BX795 reduced PDPK1 phosphorylation at Ser^241^ (Fig. 3H). There was compensatory increase in AKT^308^ + PDPK1^241^ phosphorylation and MEK phosphorylation with BRAF V600E inhibition alone and PDPK1 inhibition alone, respectively, but not with combined PDPK1 and BRAF V600E inhibition, confirming dual pathway blockade (Fig. 3H).

In addition, combination PDPK1 and BRAF V600E inhibition significantly increased apoptosis compared to either agent along and the vehicle control, with increased cleaved CASP3, PARP-1 cleavage, decreased BCL-2, and enhanced AnnexinV/PI staining (Figs. 3H–I and S3A). Furthermore, combination treatment reduced BAD phosphorylation and increased the active BAD/inactive pBAD ratio by 3.4-fold and 2.4-fold in the 8505C and 2 SW1736 ATC cell lines, respectively (Fig. 3J). Immunoprecipitation analysis revealing increased BCL-xL–BAD binding, indicating BAD-mediated inhibition of BCL-xL’s antiapoptotic function (Fig. 3K) with no binding to BCL2. Furthermore, BX795 treatment reduced Ser^9^ phosphorylation (inactive) and increased active GSK-3β levels, which were further enhanced with combination treatment (Fig. 3J). In addition, combination PDPK1 and BRAF V600E inhibition increased γ-H2AX (Ser^139^), indicating greater DNA damage than with either agent alone and the vehicle control (Fig. 3J). Overall, supervised cluster analysis demonstrated treatment-specific differential abundance across extrinsic death receptor components (TNFRSF10A and FAS), mitochondrial apoptosis regulators (BAD, BID, BAX, BAK1, and BCL2L1), and executioners of apoptosis (CASP3, CASP7, CASP8, and CASP9), with a distinct expression signature including BAD, BAX, BCL2, CASP3, and PARP1 observed with combination treatment (Fig. 3L). Combination treatment led to higher levels of BID, CASP3, and BAD and lower levels of BCL2L1 and PARP1. Although the total BAX and BCL2 protein levels were lower, the BAX/BCL2 ratio was high after combination treatment compared to either agent alone, further supporting enhanced apoptosis (Fig. 3L), Together, these results show that combination treatment synergistically induces apoptosis, activates caspase, leads to DNA damage, and suppresses survival signaling pathways in *BRAF* V600E–mutant ATC cells.

### Combination BX795 and Dab treatment induces DNA damage and G2 cell cycle arrest in ATC cells

To assess DNA damage induced by combination treatment, we evaluated the formation of γH2AX (Ser139) foci. Both BX795 and Dab treatment alone increased γH2AX, with combination treatment resulting in the highest number of foci, γH2AX-positive cells, and fluorescence intensity (Fig. 3M). Furthermore, DNA damage response activation was highest with combination treatment, with a greater increase in ATM (Ser^1981^) and CHK2 (T^68^) phosphorylation, while CHK1 (Ser^345^) phosphorylation increased with Dab treatment, decreased with BX795 treatment, and had an intermediate effect with combination treatment (Fig. 3N).

Cell cycle analysis showed BX795 induced strong G2/M arrest, Dab caused G0/G1 arrest, and combination treatment significantly increased G2/M arrest with 4N DNA accumulation (Fig. 3O). BX795 increased cyclin B, which was further elevated with combination treatment, while both agents alone reduced the CDK1/pCDK1 ratio, PLK1 phosphorylation, and phospho-CDC25C (Thr^48^), with the strongest effect observed with combination treatment (Fig. 3P). Combination treatment also lowered cyclin A2 and depleted phospho-histone H3 (Ser^10^), and disrupted spindle microtubule assembly in nocodazole-synchronized cells, indicating failure to progress from the G2 phase to the M phase despite cyclin B accumulation. Likewise, Dab treatment reduced cyclin D1, which was further reduced with combination treatment due to proteasomal degradation; however, BX795 treatment alone had no effect (Fig. 3Q). Together, these data demonstrate that combined PDPK1 and BRAF inhibition synergistically disrupts cell-cycle progression by enforcing G2/M arrest with defective mitotic entry, accompanied by proteasomal degradation of cyclin D1.

We further assessed DNA damage responses associated with combination treatment–induced G2 arrest based on the phosphoproteomic results. Combination treatment resulted in the highest phosphorylation of RAD50, CDK1, MCM2, PRKDC, and XRCC5 compared with either agent alone (Fig. 3R), consistent with enhanced sensing of double-stranded DNA (dsDNA) breaks, replication stress, reinforced G2/M checkpoint activation, and increased engagement of non-homologous end joining (NHEJ)-mediated repair. In contrast, combination treatment reduced the abundance of LIG1, MDC1, and MSH6 relative to treatment with either agent alone, indicating suppression of DNA repair synthesis, chromatin-based damage signaling, and mismatch repair pathways (Fig. 3R). Together, these findings indicate that combination treatment leads to a greater DNA damage burden marked by replication stress, checkpoint activation, and NHEJ engagement, while suppressing key DNA repair scaffolding and mismatch repair components.

Taken together, these findings suggest that dual PDPK1 and BRAF V600E inhibition activates ATM/CHK2 signaling, downregulates G2/M regulators, blocks mitotic entry, and enforces persistent G2 arrest in ATC cells.

### Combination BX795 and Dab treatment induces mitotic G2 arrest that is associated with mitochondrial dysfunction

Given that prolonged cell cycle arrest can result in oxidative stress, we assessed ROS levels following treatment with BX795, Dab, or their combination in *BRAF* V600E–mutant ATC cells. Combination treatment markedly increased intracellular ROS (DCFH-DA) and mitochondrial superoxide (MitoSOX) compared with either agent alone or the vehicle control (Fig. 4A–B). MitoTracker Green staining revealed increased mitochondrial content that was most pronounced with combination treatment (Fig. 4C–D), accompanied by a higher mitochondrial membrane potential (TMRM) without loss of integrity (Fig. 4E).

**Fig. 4:**
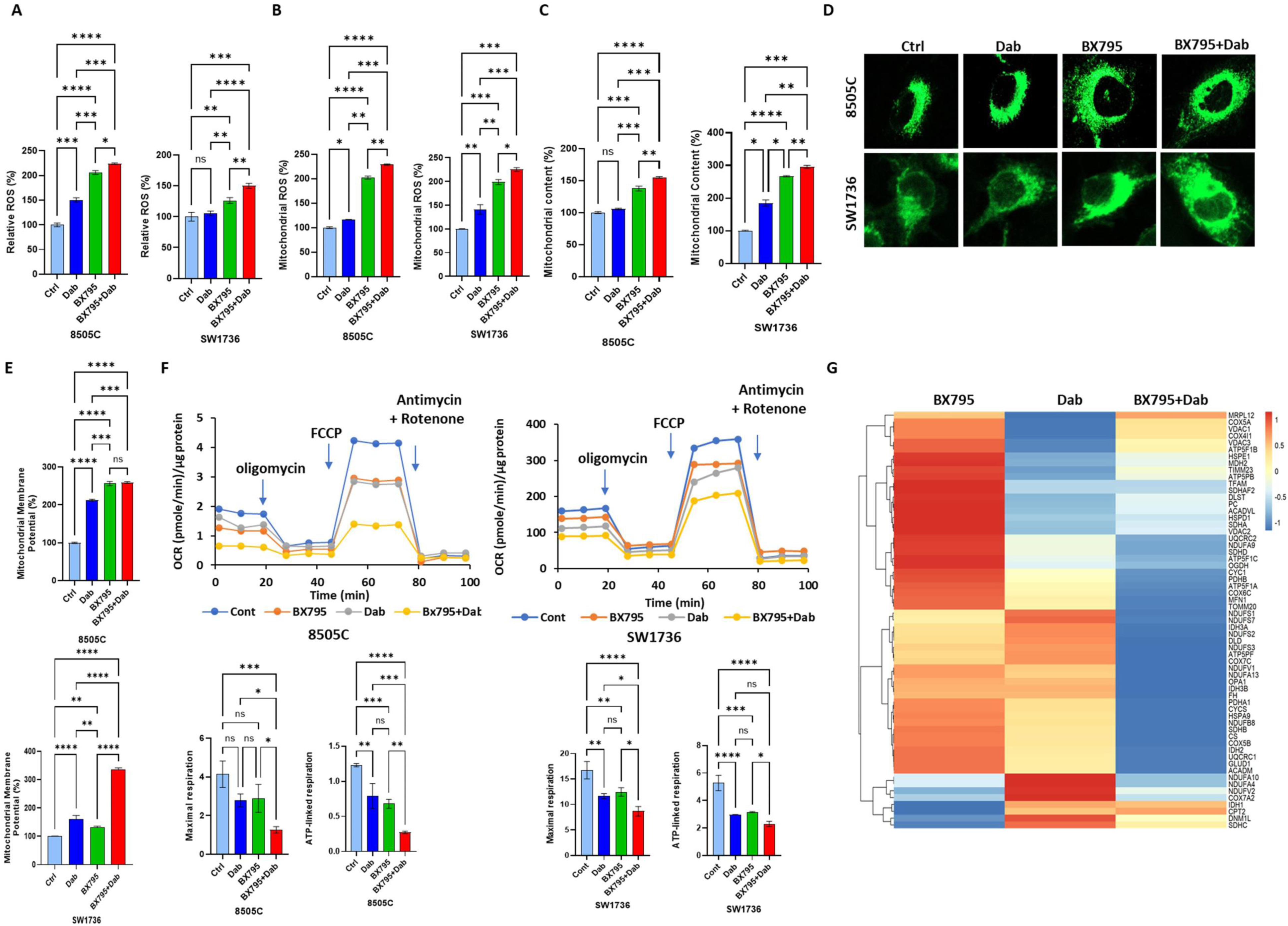
The effects of BX795 and dabrafenib (Dab) treatment on reactive oxygen species (ROS), mitochondrial function, and bioenergetics in *BRAF* V600E–mutant anaplastic thyroid cancer (ATC) cell lines. Mutated BRAF V600E cell were treated with BX795 (2.5 µM), Dab (2.5 µM), or their combination. (A) Total cellular ROS was measured using the DCFH-DA fluorescent dye and quantified with flow cytometry. (B) Mitochondrial superoxide was measured based on MitoSOX red fluorescence, quantified with flow cytometry. (C–D) The mitochondrial content was assessed by MitoTracker green fluorescence, as quantified by flow cytometry and visualized with live-cell confocal microscopy (Zeiss LSM 800, 400× magnification). (E) The mitochondrial membrane potential was evaluated by TMRM (100 nM) fluorescence and analyzed by flow cytometry. (F) The oxygen consumption rate (OCR) was measured using a Seahorse analyzer with sequential injections of oligomycin (Oligo), 2,4-dinitrophenol (DNP), and antimycin A plus rotenone (Rot+AA). The data were normalized to the total protein content, and maximal respiration and adenosine triphosphate (ATP)-linked respiration were quantified accordingly. (G) The heatmap shows z-scored log_2_-transformed abundance of proteins associated with mitochondrial oxidative phosphorylation, the tricarboxylic acid (TCA) cycle, mitochondrial ribosomal, transport, and mitochondrial dynamics–associated proteins in 8505C ATC cells treated with BX795 (2.5 µM), Dab (2.5 µM), or their combination. The proteins include representatives of electron transport chain complexes I–V (NDUF subunits, SDHA–D, UQCRC1/2, COX subunits, and ATP5 subunits), TCA cycle enzymes (IDH1, IDH2, IDH3, FH, OGDH, DLST, CS, MDH2, and PDHA1/PDHB), mitochondrial transport and structure (TOMM20, TIMM23, VDAC1–3, MFN1, OPA1, DNM1L), and mitochondrial genome maintenance (TFAM). Hierarchical clustering was applied to the rows. The color scale represents relative protein abundance (red = higher; blue = lower). All data are expressed as the mean ± standard deviation. Statistical significance is indicated as ns = nonsignificant; *p < 0.05; **p < 0.01; ***p < 0.001; ****p < 0.0001.

Seahorse analysis showed that combination treatment significantly reduced basal, ATP-linked, and maximal respiration, indicating impairment in oxidative phosphorylation (Figs. 4F and S4A). Mitochondrial integrity was also confirmed by JC-1 staining (Fig. S4B). Profiling of proteins related to oxidative phosphorylation revealed complex, specific changes with combined treatments, where complex V increased in both *BRAF* V600E–mutant ATC cell lines. On the other hand, relative to the vehicle control, complex III was higher in 8505C cells, while complex II and IV were higher in SW1736 cells (Fig. S4C).

We performed supervised cluster analysis of the proteomic data to determine the effect of BX795, Dab, and their combination on proteins involved in mitochondrial ribosomal function, electron transport chain complexes, the tricarboxylic acid (TCA) cycle, mitochondrial dynamics, and metabolite transport (Fig. 5G). We found higher relative abundance of mitochondrial respiratory chain and metabolic enzymes with BX795 treatment, including multiple complex I subunits (NDUFS1, NDUFS7, NDUFS2, NDUFS3, NDUFV1, and NDUFB8), complex II components (SDHA, SDHB, SDHC, and SDHD), complex III proteins (UQCRC1 and UQCRC2), complex IV subunits (COX5A, COX4I1, COX6C, COX7C, COX5B, and COX7A2), and ATP synthase subunits (ATP5F1A, ATP5F1B, ATP5F1C, ATP5PB, and ATP5PF). There was reduced relative abundance with Dab treatment and markedly reduced abundance with combination treatment. Proteins involved in the TCA cycle and mitochondrial metabolism, including IDH1, IDH2, IDH3A, IDH3B, FH, OGDH, DLST, MDH2, CS, PDHA1, PDHB, PC, DLD, GLUD1, ACADM, and ACADVL, followed a similar pattern, with higher expression with BX795 treatment, intermediate or reduced expression with Dab treatment, and the lowest relative abundance with combination treatment.

**Fig. 5:**
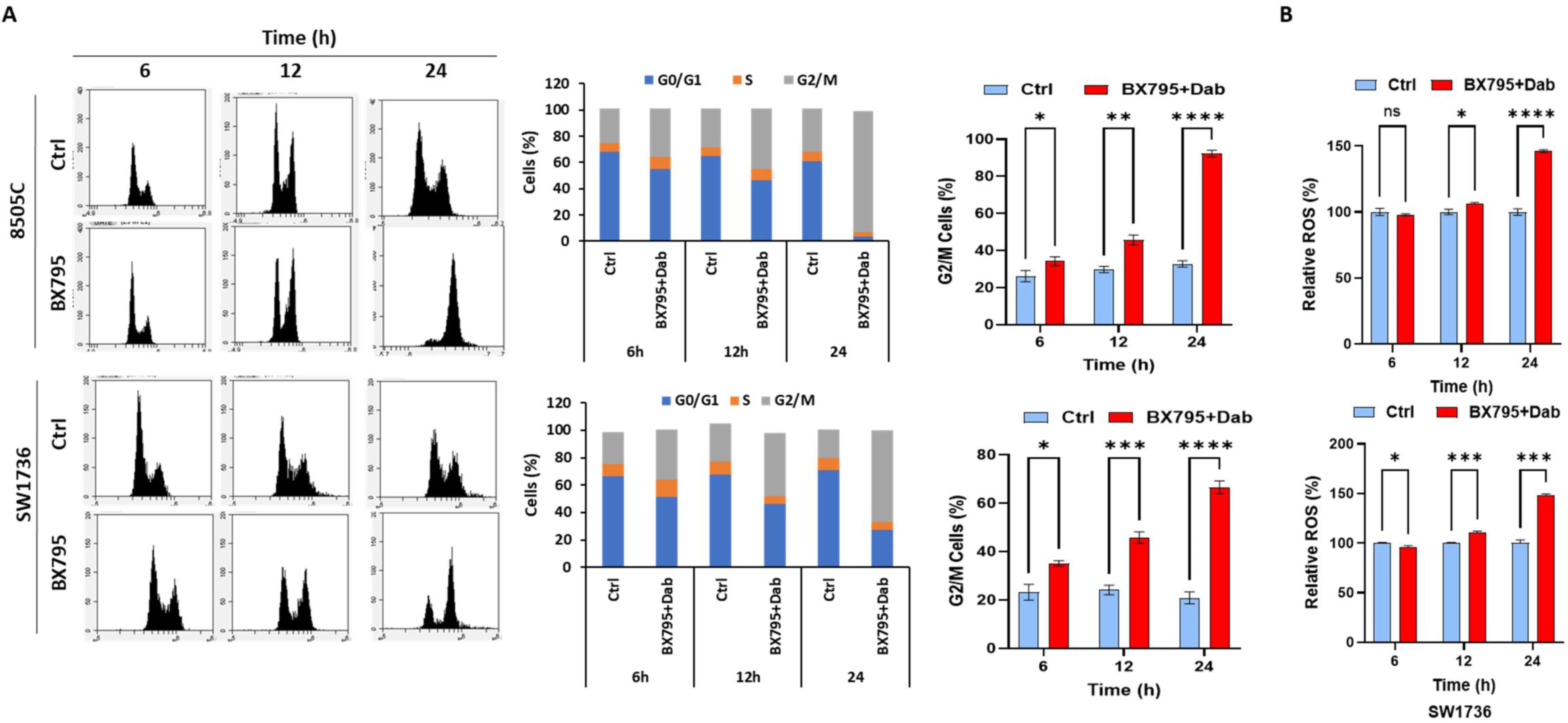
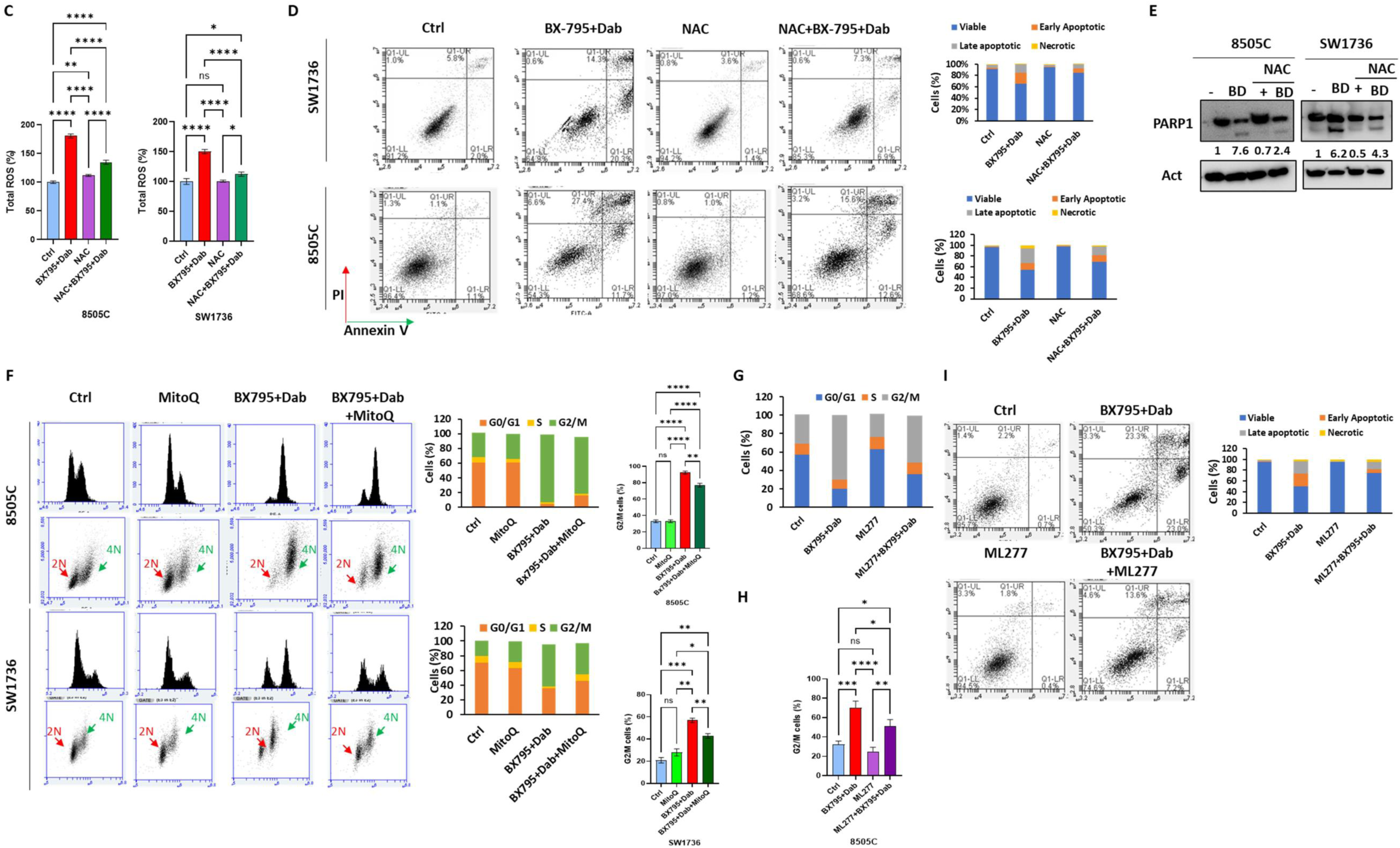
Reactive oxygen species (ROS) scavenging attenuates combination BX795 and dabrafenib (Dab) treatment effect while CHK2 inhibition leads to mitotic slippage and partial apoptotic reversal. (A) Time-dependent effect of combination BX795 and Dab treatment based on propidium iodide (PI) staining, showing the proportion of cells arrested in the G2/M phase. Cells were stained with PI and analyzed by fluorescence-activated cell sorting. The results are presented as the mean ± standard deviation. Statistical significance is indicated as ns = nonsignificant; *p < 0.05; **p < 0.01; ***p < 0.001; ****p < 0.0001. (B) Intracellular ROS levels were measured using DCFH-DA staining and analyzed by flow cytometry. The results are presented as the mean ± standard deviation. (C–E) The effects of *N*-acetyl cysteine (NAC) (5 mM) on combination treatment–induced responses in *BRAF* V600E–mutant ATC cell lines. (C) ROS generation was analyzed by DCFH-DA staining and is presented as the mean ± standard deviation. (D) Apoptosis was assessed based on Annexin V/PI staining. (E) Immunoblot analysis of PARP-1 cleavage using β-actin as a loading control. The densitometric quantification of cleaved vs pro-PARP cleavage in relation to loading control is shown below the bands. (F) The effect of the mitochondrial ROS scavenger MitoQ on combination treatment–induced G2/M arrest in *BRAF* V600E–mutant ATC cell lines. The DNA content was measured, 2N (diploid) and 4N (tetraploid), based on PI staining. The results are presented as the mean ± standard deviation. Statistical significance is indicated as ns = nonsignificant; *p < 0.05; **p < 0.01; ***p < 0.001; ****p < 0.0001. (G–H) The impact of CHK2 phosphorylation inhibitor BML-277 (ML277) on combination BX795 and Dab treatment-induced. (G) Cell cycle arrest was analyzed by PI staining and flow cytometry. (H) Antiproliferative effect in *BRAF* V600E–mutant ATC cells. The DNA content was measured, 2N (diploid) and 4N (tetraploid), based on PI staining. The results are presented as the mean ± standard deviation. Statistical significance is indicated as ns = nonsignificant; *P < 0.05; **P < 0.01; ***P < 0.001; ****P < 0.0001. (I) Apoptosis was measured with Annexin V/PI staining and flow cytometry. The results are presented as mean ± standard deviation. Statistical significance is indicated as ns = nonsignificant; *p < 0.05; **p < 0.01; ***p < 0.001; ****p < 0.0001.

Mitochondrial structure, dynamics, and transport proteins, including TFAM, TOMM20, TIMM23, MFN1, OPA1, DNM1L, VDAC1, VDAC2, and VDAC3, also a demonstrated progressive reduction from BX795 to Dab treatment, and a further reduction with combination treatment. CYCS and HSPA9 displayed higher abundance with BX795 treatment, reduced abundance with Dab treatment, and further reduction with combination treatment. Collectively, these data demonstrate broad and coordinated suppression of mitochondrial oxidative phosphorylation, TCA cycle activity, and mitochondrial structural components with combination treatment, with intermediate effects observed with Dab treatment alone and the highest relative expression with BX795 treatment.

The consistent decrease in mitochondrial complex proteins supports that mitochondrial dysfunction occurs with dual PDPK1 and BRAF V600E inhibition. This triggers ROS generation even though the mitochondrial membrane potential is preserved. These changes are consistent with a glycolytic, senescence-like phenotype as described previously (47).

### ROS scavenging mitigates genotoxicity while CHK2 inhibition induces mitotic slippage and partially reverses combination therapy-induced apoptosis

Next, we preformed time-course analysis to determine the sequence of events with combination treatment that leads to its synergistic anticancer activity. We found G2/M arrest occurred by 6 h; this preceded ROS elevation (which began at 12 h) (Fig. 5A–B). The mitochondrial membrane potential increased as early as 6 h, while mitochondrial accumulation occurred later (≥ 20 h) (Fig. S4D–E). Thus, combination treatment induces early G2/M arrest and mitochondrial hyperpolarization, followed by sustained ROS production in *BRAF* V600E–mutant ATC cells. To assess the role of ROS in combination treatment–induced cytotoxicity, we used the ROS scavengers *N*-acetyl cysteine (NAC) and MitoQ. NAC significantly reduced combination treatment–induced ROS (DCFH-DA) (Fig. 5C), apoptosis (Annexin V/PI) (Fig. 5D), and PARP cleavage (Fig. 5E), indicating ROS-mediated apoptosis. NAC treatment also decreased γ-H_2_AX and DNA damage response proteins (pATM and pCHK2) but did not reverse G2 arrest (Fig. S4F), suggesting ROS generation occurs downstream of cell cycle arrest. MitoQ treatment partially released cells from G2 arrest (Fig. 5F), implicating mitochondrial superoxide in cell cycle regulation.

Next, to explore the involvement of CHKs in combination treatment–induced cell cycle arrest, we used the CHK2 inhibitor BML-277, which significantly reversed combination treatment–induced G2 arrest (Figs. 5G and S4G), cell proliferation (Fig. 5H), and apoptosis (Fig. 5I) in 8505C cells. As CHK1 and CHK2 both were active in SW1736 cells, we used AZD7762 (which inhibits both CHK1 and CHK2) in SW1736 cells, which resulted in mitotic slippage (Fig. S4H) and reversed the inhibition of cellular proliferation (Fig. S4I). These data suggest that mitotic arrest leads to apoptosis, which can be partially reversed by mitotic slippage.

## DISCUSSION

PDPK1 is overexpressed in thyroid cancer (39). High PDPK1 levels in thyroid cancer are associated with advanced stage, larger tumor size, lymph node metastasis, ATC, and worse overall and recurrence-free survival (48). In this study, we found high levels of total and phosphorylated PDPK1 at Ser241 in ATC cells, and combined inhibition of PDPK1 and BRAF V600E had synergistic anticancer activity in *BRAF* V600E–mutant ATC in *in vitro*, *ex vivo*, and *in vivo* models. Mechanistically, this combination effectively suppressed both the PDPK1/AKT and MAPK pathways without rebound activation that is seen with inhibition of a single pathway. Dual inhibition resulted in pronounced DNA damage, G2 cell cycle arrest, and apoptosis in *BRAF* V600E–mutant ATC cells.

Dual inhibition of PDPK1 and mutant *BRAF* V600E–mutant ATC drives accumulation of dsDNA damage and G2 arrest through complementary mechanisms, where *BRAF* V600E inhibition induces replication stress by abruptly halting MAPK-driven proliferation, generating stalled replication forks that can collapse into dsDNA breaks (49). PDPK1 blockade suppresses AKT and other AGC kinase signaling, impairing key DNA repair pathways including homologous recombination and NHEJ, thereby preventing resolution of these molecular events (50,51). The resulting accumulation of unrepaired DNA damage activates the G2/M checkpoint, enforcing cell cycle arrest and sensitizing cells to apoptosis (52). We found the combined inhibition of PDPK1 and *BRAF* V600E results in accumulation of unrepaired DNA damage, which activates the G2/M checkpoint and enhances apoptotic commitment, as observed in our cell viability and apoptosis assays. Interestingly, this genotoxic stress is also accompanied by mitochondrial dysfunction, characterized by disrupted oxidative phosphorylation and an unexpected increase in the mitochondrial membrane potential, without triggering mitochondrial outer membrane permeabilization. This hyperpolarized mitochondrial state likely reflects an adaptive response to maintain ATP production under DNA damage–induced metabolic stress, consistent with previous observations that DNA damage can alter mitochondrial bioenergetics and promote mitochondrial hyperpolarization (53,54). This mechanistic interplay aligns with studies linking cell cycle checkpoint disruption to mitochondrial dysfunction, ROS accumulation, and apoptotic priming (55).

Treatment with the mitochondria-targeted antioxidant MitoQ partially reduced G2 arrest, suggesting that mitochondrial ROS and hyperpolarization contribute to the enforcement of cell cycle checkpoints. This is consistent with reports that mitochondrial ROS can act as signaling intermediates to activate DNA damage checkpoints and maintain G2 arrest (53,56). These findings imply that mitochondrial bioenergetic stress is not merely a byproduct of DNA damage but actively reinforces G2/M checkpoint signaling. Therapeutically, this highlights that combined inhibition of PDPK1 and BRAF V600E exploits a dual vulnerability: suppression of proliferative signaling and engagement of DNA damage responses. These results highlight a feed-forward mechanism where mitochondrial dysfunction amplifies cell cycle arrest, providing a therapeutic vulnerability that is exploitable by combined inhibition of PDPK1 and BRAF V600E.

Dual inhibition of PDPK1 and *BRAF* V600E promoted dsDNA breaks and oxidative stress. Notably, treatment with the ROS scavenger NAC attenuated apoptosis and reduced DNA damage but did not reverse G2 arrest, indicating that mitochondrial ROS primarily function downstream of cell cycle checkpoint activation to drive cell death rather than initiating the arrest itself. NAC primarily scavenges H_2_O_2_, HOCl, and •OH, increasing GSH levels (57), whereas MitoQ targets mitochondrial superoxide and H_2_O_2_ (58). Because ROS species can propagate via chain reactions (59,60), MitoQ-mediated arrest release supports a role for mitochondrial superoxide in sustaining G2 arrest during combination treatment. These findings suggest a model in which DNA damage response–mediated G2 arrest primes mitochondria toward a hyperpolarized, pro-apoptotic state, with oxidative stress acting as a critical amplifier of apoptosis in ATC cells (53,56). Although the OCR was not reduced, mitochondrial hyperpolarization reflects impaired proton dissipation and electron congestion within the electron transport chain rather than enhanced oxidative phosphorylation, a characteristic feature of dysfunctional, ROS-producing mitochondria (61). BAD phosphorylation by AKT and RSK1—regulated by PDPK1 and MAPK—leads to inactivation and sequestration by 14-3-3 proteins, a mechanism linked to BRAF inhibitor resistance in melanoma (62). Inhibiting PI3K/PDPK1/AKT and BRAF pathways prevents BAD sequestration and promotes active BAD driven apoptosis (63–65). This mechanism underscores the therapeutic potential of combining PDPK1 and BRAF V600E inhibition to simultaneously enforce cell cycle blockade and trigger ROS-dependent apoptotic pathways.

ATC harbors frequent mutations in genes that lead to activation of both the PI3K/AKT/mTOR and MAPK signaling pathways. Activating mutations in PIK3CA occur in approximately 10%–20% of ATC cases, while PTEN loss or inactivating mutations are also common, collectively promoting constitutive PI3K/AKT signaling (4,66,67). In parallel, MAPK pathway activation is highly prevalent, with *BRAF* V600E mutations detected in 40%–70% of ATC cases, and RAS mutations contributing to MAPK hyperactivation in a significant subset (4,68). These two pathways are activated in ATC, reinforcing proliferative, survival, and stress-adaptive signaling. Their combined deregulation underpins therapeutic resistance to *BRAF* V600E inhibitors and a poor prognosis in ATC (63,69,70), underscoring the need for rational dual-targeting strategies. PDPK1 represents a strategic therapeutic node in ATC because it lies downstream of PI3K and integrates signals from multiple recurrent alterations, including *PIK3CA*, *PTEN*, and *RAS*. Selective inhibition of PI3K or AKT may leave bypass pathways intact. On the other hand, PDPK1 activates a broad network of AGC kinases (AKT, SGK, S6K, and PKC isoforms), making it a more effective point of signaling blockade. By targeting PDPK1, both survival signaling and stress-adaptation programs are disrupted, addressing the multifaceted oncogenic landscape of ATC.

The therapeutic implications of our findings are immediate. Dab is a Food and Drug Administration–approved inhibitor of BRAF V600E, while the PDPK1 inhibitor BX795 has antitumor activity in *in vitro*, *ex vivo*, and *in vivo* models of ATC and resulted in no treatment-related toxicity alone or in combination with Dab *in vivo*. Our data show that PDPK1 inhibition exerts intrinsic anticancer activity and significantly enhances BRAF V600E inhibition, supporting a rational, dual-targeted approach. Given that both compounds have been validated preclinically and/or clinically, this combination represents an opportunity to translate this combination into an early-phase clinical trial in *BRAF* V600E–mutant ATC. Moreover, because *BRAF* V600E is present in ∼15% of all human cancers, this strategy may extend beyond ATC to other malignancies driven by MAPK and PI3K/AKT/mTOR crosstalk.

In summary, we have identified PDPK1 as a critical therapeutic vulnerability in ATC. By functioning as a central effector of PI3K/AKT/mTOR signaling, PDPK1 integrates multiple oncogenic inputs common in ATC, and its inhibition disrupts both survival and stress-adaptation pathways. Importantly, combining PDPK1 inhibition with BRAF blockade yields synergistic anticancer activity. These findings provide a strong rationale for advancing PDPK1 inhibition, particularly in combination with *BRAF* V600E inhibition, into the clinic.

## Author Contribution declaration

Tejinder Pal Khaket: Conceptualization, Methodology, Validation, Investigation, Visualization, Writing- Original draft preparation. Chandrayee Ghosh, Zhongyue Yang, Jiangnan Hu, Eden D. Alamaw, Myriem Boufraqech, Thorkell Andresson Maura O’Neill, Ya-Qin Zhang, Min Shen, Bereketeab Haileselassie: Visualization, Investigation, Data curation, Methodology. Electron Kebebew: Supervision, Conceptualization, Project administration, Writing- Reviewing and Editing, Funding acquisition.

## Consent to Publish declaration

Not applicable

## Ethics Approval and Consent to Participate

Patient-derived tumor samples were obtained under protocols approved by the Stanford University Institutional Review Board (IRB protocol 50782). All samples were collected with written informed consent from participants in accordance with institutional guidelines. No identifying information was used.

All animal procedures were approved by the Stanford Administrative Panel on Laboratory Animal Care (APLAC; Protocol #34767) and conducted in accordance with institutional and federal guidelines.

## Availability of data and materials

All data generated or analyzed during this study are presented within the manuscript and its supplementary materials. Any additional information or raw datasets can be obtained from the corresponding author upon reasonable request.

## Supplemental Figures

**Fig. S1: PDPK1 expression, the efficacy of PDPK1 inhibitors, and the effect of combination PDPK1 and BRAF V600E inhibition in anaplastic thyroid cancer (ATC) cells.**

(A) The box-and-whisker plot shows the Chronos gene effect scores for PDPK1 across the top 10 cancer lineages from the DepMap 25Q3 dataset. The scores represent the impact of PDPK1 knockout on cell fitness, where more negative values indicate greater essentiality. Individual dots represent cell lines within each lineage, and boxes denote the median and interquartile range. The thyroid cancer lineage is highlighted in red. (B) PDPK1 messenger RNA (mRNA) expression presented as Expression log_2_(TPM+1) among different thyroid cancer cell lines: ATC, follicular thyroid cancer (FTC), well-differentiated thyroid cancer (WDTC), medullary thyroid cancer (MTC), and poorly differentiated thyroid cancer (PDTC). (C) The box-and-whicker plot shows the Chronos gene effect scores for PDPK1 across thyroid cancer subtypes from the DepMap 25Q3 dataset. Individual data points represent cell lines, and boxes denote the median and interquartile range. ATC is highlighted in red. (D) Data from quantitative high-throughput drug screening in ATC cell lines (8505C, THJ16T, and THJ11T). Comparison of half-maximal inhibitory concentration (IC_50_) < 10 μM and drug-response curve class of < 2.2 for known PDPK1 inhibitors (BX795, GSK2334470, OSU-03012, DTXSID90678515, and BX912) after treatment for 48 h. (E) PRISM drug sensitivity scores for BX795 (24Q2) across thyroid cancer subtypes, including WDTC, ATC, PDTC, and MTC. More negative values indicate greater sensitivity to BX795. The data are derived from pooled cell line responses within each subtype. (F) To analyze the effect of BX795 treatment, *BRAF* V600E–mutant ATC cell lines (8505C and SW1736) were treated with BX795 (2.5 or 5 µM) for 48 h, and then the levels of non-phosphorylated and phosphorylated mTOR, AKT (T^308^), H_2_AX (S^15^), and S6K were evaluated by immunoblotting using β-actin as a loading control. (G) *BRAF* V600E–mutant ATC cell lines (8505C and SW1736) were transiently silenced with Control siRNA, PDPK1si1, or PDPK1si2 in combination with BX795 (2.5 µM) for 48 h, and then the levels of phosphorylated PDPK1 (S^245^), H_2_AX, and PDPK1 were evaluated by immunoblotting using β-actin as a loading control. Their effect on cell proliferation was also measured. The results are presented as the mean ± SD. Statistical significance is indicated as ns = nonsignificant; *p < 0.05; **p < 0.01; ***p < 0.001; ****p < 0.0001. (H–I) Representative images of the colony-formation assay (H) and cellular migration (I) with BX795 and dabrafenib (Dab) treatment alone and their combination in *BRAF* V600E–mutant ATC cells. The representative images correspond to the data shown in Fig. 1K–N.

**Fig. S2: Combination BX795 and dabrafenib (Dab) treatment *in vivo*.**

(A–B) Representative tumor images from all mouse groups at the end of the study by sex. (C–D) Analysis of the tumor weight (C) and tumor volume (D) at the study endpoint by sex. The results are presented as the mean ± standard error of the mean. Statistical significance is indicated as ns = nonsignificant; *p < 0.05.

**Fig. S3: Combination BX795 and dabrafenib (Dab) treatment induces apoptosis and G2 arrest.**

(A) The effect of treatment with BX795 (2.5 µM), Dab (2.5 µM), or their combination for 40 h on BCL2 and phospho-PDPK1 (pPDPK1) protein levels in *BRAF* V600E–mutant ATC cells (8505C). Immunofluorescence was performed with BCL2 labeled using Alexa Fluor™ 633 secondary antibody and pPDPK1 labeled with Alexa Fluor™ 546 secondary antibody; DAPI was used as a nuclear counterstain. Images were acquired using a Zeiss LSM 800 confocal microscope at 400× magnification. (B) Densitometry results for the proteins shown in Fig. 4P, normalized to β-actin. The y-axis is the fold change relative to the control.

**Fig. S4: The effect of combination BX795 and dabrafenib (Dab) treatment on basal respiration, the mitochondrial membrane potential, DNA damage proteins, and cell cycle arrest.**

(A) Analysis of the effect of treatment with BX795, Dab, or their combination on basal respiration in *BRAF* V600E–mutant ATC cell lines (8505C and SW1736), measured with the Seahorse assay. The basal respiration rate was normalized to the total protein content and is presented as the mean ± standard deviation. (B) Mitochondrial integrity was assessed by JC-1 staining after combination treatment, with the red-to-green fluorescence ratio determined with flow cytometry. The results are presented as mean ± standard deviation. (C) Immunoblot analysis of oxidative phosphorylation proteins following treatments, using β-actin as a loading control. Protein band density was determined and normalized to β-actin. The y-axis is the fold change relative to the control (DMSO). (D–E) Time-dependent effects (0–40 h) of treatment with BX795 (2.5 µM), Dab (2.5 µM), or their combination on the mitochondrial membrane potential (D) and mitochondrial content (E) in *BRAF* V600E–mutant ATC cell lines. The mitochondrial membrane potential was assessed using TMRM (100 nM) fluorescence, and the mitochondrial content was measured with MitoTracker Green dye, with flow cytometry used for quantification. The data are presented as the mean ± standard deviation. (F) Immunoblot analysis of phosphorylated CHK2 (pCHK2), phosphorylated ATM (pATM), and phosphorylated H2AX (pH2AX) levels in *BRAF* V600E–mutant ATC cell lines treated with BX795 (2.5 µM) and Dab (2.5 µM) in the presence or absence of *N*-acetyl cysteine (NAC) (5 mM). β-Actin was used as a loading control. (G) Representative results of Fig. 5G. (H–I) Analysis of the effect of the AZD7762 on cell cycle arrest and the antiproliferative effect induced by combination BX795 and Dab treatment in *BRAF* V600E–mutant ATC cells (SW1736). The results are presented as the mean ± standard deviation. Statistical significance is indicated as ns = nonsignificant; *p < 0.05; **p < 0.01; ***p < 0.001; ****p < 0.0001.

