## supplementary tables for "Dual targeting of PDPK1 and BRAF V600E is synthetically lethal": supplementary tables.pdf

**Table S1. List of antibodies.**

| Antibodies | Product ID | Manufacturer |
|---|---|---|
| $pPDPK1^{s231}$ | 3061 | CST |
| PDPK1 | 5662 | CST |
| pMEK1/2 | 2338 | CST |
| MEK1/2 | 8727 | CST |
| $pAKT^{308}$ | 9275 | CST |
| AKT | 4691 | CST |
| PARP-1 | 9542 | CST |
| BCL2 | 50071 | CST |
| Caspase-3 | 9661 | CST |
| $pBAD^{S112}$ | 5284 | CST |
| BAD | 9292 | CST |
| BCL-xl | 2764 | CST |
| $pGSK3\beta^{Ser9}$ | 9323 | CST |
| GSK3β | 12456 | CST |
| $pATM^{S1981}$ | Ab81292 | Abcam |
| ATM | 2873 | CST |
| $pCHK2^{T68}$ | 2197 | CST |
| CHK2 | 2662 | CST |
| $pCHK1^{S345}$ | 2348 | CST |
| Cyclin D | 2978 | CST |
| Cyclin B | 12231 | CST |
| $pcdc25C^{T48}$ | 4901 | CST |
| $pCDK1^{Y15}$ | 4539 | CST |
| CDK1 | 9116 | CST |
| $pH3A^{S10}$ | 3377 | CST |
| H3A | PAS-16183 | Invitrogen |
| Cyclin A2 | 4656 | CST |
| $pH2AX^{S139}$ | 2577 | CST |

| | | |
|---|---|---|
| H2AX | 2595 | CST |
| OXPhos | PK30006 | Proteintech |
| α-Tubulin | 3873 | CST |
| Alexa Fluor™ 488 | A11001 | Invitrogen |
| Alexa Fluor™ 546 | A11035 | Invitrogen |
| Alexa Fluor™ 633 | 21052 | Invitrogen |
| β-actin | 3700 | CST |
| Anti-rabbit IgG, HRP-linked antibody #7074 | 7074 | CST |
| Anti-mouse IgG, HRP-linked antibody #7076 | 7074 | CST |

**Table S2. List of chemicals.**

| **Chemicals** | **Product ID** | **Source** |
|---|---|---|
| BX795 | S1274 | Selleckchem |
| Dabrafenib | S2807 | Selleckchem |
| BML-277 | S8632 | Selleckchem |
| NAC | A15409.36 | Thermo Fisher Scientific |
| MitoQ | S8978 | Selleckchem |
| Oligomycin | 75351 | Sigma-Aldrich |
| FCCP | S8276 | Selleckchem |
| Antimycin | A8674 | Sigma-Aldrich |
| 0.25% Trypsin-EDTA | 25200056 | Thermo Fisher Scientific |
| Western Blot Stripping Buffer | T7135A | Takara Bio Inc. |
| 4× Laemmli Sample Buffer | 1610747 | Bio-Rad |
| Lipofectamine™ 2000 Transfection Reagent | 11668027 | Thermo Fisher Scientific |
| Opti-MEM™ I Reduced Serum Medium | 31985070 | Thermo Fisher Scientific |
| Phosphatase and protease inhibitors | PI78441 | Thermo Fisher Scientific |
| NP-40 Surfact-Amps™ Detergent Solution | 85124 | Thermo Fisher Scientific |
| ProLong™ Gold Antifade Mountant | P10144 | Thermo Fisher Scientific |
| DMEM/F-12 | 21331020 | Thermo Fisher Scientific |
| Protein A/G Plus -Agarose | Sc-2003 | Santa Cruz |
| RIPA Lysis Buffer | 89901 | Thermo Fisher Scientific |
| Sodium Bicarbonate 7.5% solution | 25080094 | Thermo Fisher Scientific |
| Sodium Pyruvate (100 mM) | 11360070 | Thermo Fisher Scientific |
| Invitrogen™ UltraPure™ DNase/RNase-Free Distilled Water | 10-977-015 | Thermo Fisher Scientific |
| MitoSOX™ Red Mitochondrial Superoxide Indicator, for live-cell imaging | M36008 | Thermo Fisher Scientific |
| Microplates for Fluorescence-based Assays, 96-well | 3904 | Thermo Fisher Scientific |

| | | |
|---|---|---|
| Corning™ DMEM with L-Glutamine, 4.5g/L Glucose and Sodium Pyruvate | 10013CV | Fisher Scientific |
| Antibiotic Antimycotic Solution (100×) | A5955 | Sigma Aldrich |
| Seahorse XFe24 FluxPak kit | 102340-100 | Agilent |
| Bovine serum albumin | BP9703-100 | Fisher Scientific |
| Bio-Rad Protein Assay Dye | 5000006 | Bio-Rad |
| Nocodazole | 31430-18-9 | Sigma Aldrich |
| BD Pharmigen FITC Annexin V apoptosis detection Kit II | 51-6710AK | BD Bioscience |
| Bio-Rad Mini-PROTEAN TGX gels 4-15% | 4561086 | Bio-Rad |
| Formalin Solution | SF98-4 | Fisher Scientific |
| Crystal Violet 1% Aquos Solution | V5265 | Sigma Aldrich |
| Methyl-cellulose | AC428432500 | Fisher Scientific |
| CellTiter-Glo®, | G9241 | Promega |
| 3D Tumorsphere MediumXF | C-28070 | PromoCell |
| EasyPep™ lysis buffer | A45735 | Thermo Fisher Scientific |
| PhosSTOP™ | 4906845001 | Sigma Aldrich |
| FxCycleTM PI/RNase staining solution | F10797 | Thermo Fisher Scientific |

**Table S3. Thyroid cancer cell lines and their mutations.**

| Cell line | Mutation status |
|---|---|
| **SW1736** | **BRAF, p.Val600Glu (c.1799T>A), heterozygous**<br>TERT, c.1–124C>T (c.228C>T), heterozygous<br>TP53, p.Gln192Ter (c.574C>T), homozygous;<br>TSHR, p.Ile486Phe (c.1456A>T), heterozygous |
| **8505C** | **BRAF, p.Val600Glu (c.1799T>A), homozygous**<br>NF2, p.Glu129Ter (c.385G>T), homozygous<br>TERT, c.1–146C>T (c.250C>T), heterozygous<br>TP53, p.Arg248Gly (c.742C>G), hemizygous |
| **BCPAP** | **BRAF, p.Val600Glu (c.1799T>A), homozygous**<br>TERT, simple, c.1–124C>T (c.228C>T), heterozygous<br>TP53, simple, p.Asp259Tyr (c.775G>T), homozygous |
| **THJ29T** | FGFR2 + HGNC<br>CDKN2A, homozygous<br>HDAC10 (c.399delG); heterozygous<br>TERT; simple; (c.250C>T), heterozygous<br>TP53; simple; p.Gln104Ter (c.310C>T), homozygous |
| **THJ16T** | MKRN1, Name(s)=MKRN1-BRAF, HGNC, HGNC:3373, EP300, p.Ser799Phefs*5 (c.2396delC), homozygous<br>**PIK3CA, p.Glu545Lys (c.1633G>A), heterozygous**<br>RET, p.Glu90Lys (c.268G>A), heterozygous<br>TERT, c.1–124C>T (c.228C>T) (C228T), heterozygous<br>TP53, p.Arg273His (c.818G>A), homozygous |
| **THJ11T** | KRAS; simple; p.Gly12Val (c.35G>T); zygosity = unspecified<br>TERT; simple; c.1-124C>T (c.228C>T) (C228T); zygosity = heterozygous |
| **C643** | HRAS, p.Gly13Arg (c.37G>C), unspecified<br>**PTEN, p.Phe341Leu (c.1023T>G), heterozygous**<br>TERT, c.1–124C>T (c.228C>T), heterozygous<br>TP53, p.Arg248Gln (c.743G>A), homozygous |

| | |
|---|---|
| | VTCN1, p.Tyr215Ter (c.645C>G), unspecified |
| **FTC236** | MSH6, p.Lys1045fs (c.3135delG), homozygous<br>**PTEN, p.Arg130Ter (c.388C>T), homozygous**<br>RB1, p.Arg320Ter (c.958C>T), homozygous<br>TERT, c.1–124C>T (c.228C>T), homozygous<br>TP53, p.Arg273His (c.818G>A), homozygous |
| **FTC133** | FLCN, p.His429Thrfs*39 (c.1285delC), homozygous<br>MSH6, p.Lys1045fs (c.3135delG), homozygous<br>NF1, p.Cys167Ter (c.501T>A), homozygous<br>**PTEN, p.Arg130Ter (c.388C>T), homozygous**<br>TERT, c.1–124C>T (c.228C>T) (C228T), homozygous<br>TP53, p.Arg273His (c.818G>A), homozygous |
| **TPC1** | CCDC6-RET, RET/PTC1, CDKN2A, p.Ala68fs, homozygous<br>STAG2, p.Gln1089Ter (c.3265C>T), heterozygous<br>TERT, c.1–124C>T (c.228C>T), heterozygous |

**Table S4. Oncogenic driver mutations and PDPK1 dependency in thyroid cancer.**

| **Disease** | **Mutation(s)** | **Cell line** | **Chronos** |
|---|---|---|---|
| Anaplastic thyroid cancer | *BRAF* V600E | BHT101 | –1.15 |
| | | 8305C | –0.84 |
| | | 8505C | –0.781 |
| | *PTEN* F241fs | ASH3 | –0.656 |
| | *KRAS* G12R | CAL62 | –1.01 |
| Well-differentiated thyroid cancer | *AKT* E17K +*BRAF* V600E | IHH4 | –0.985 |
| | *PTEN* R130Ter | FTC133 | –0.606 |
| Poorly differentiated thyroid cancer | *BRAF* V600E | BCPAP | –0.88 |
