## Supplementary figures and images for "Dual targeting of PDPK1 and BRAF V600E is synthetically lethal"

### supplementary figures.pdf

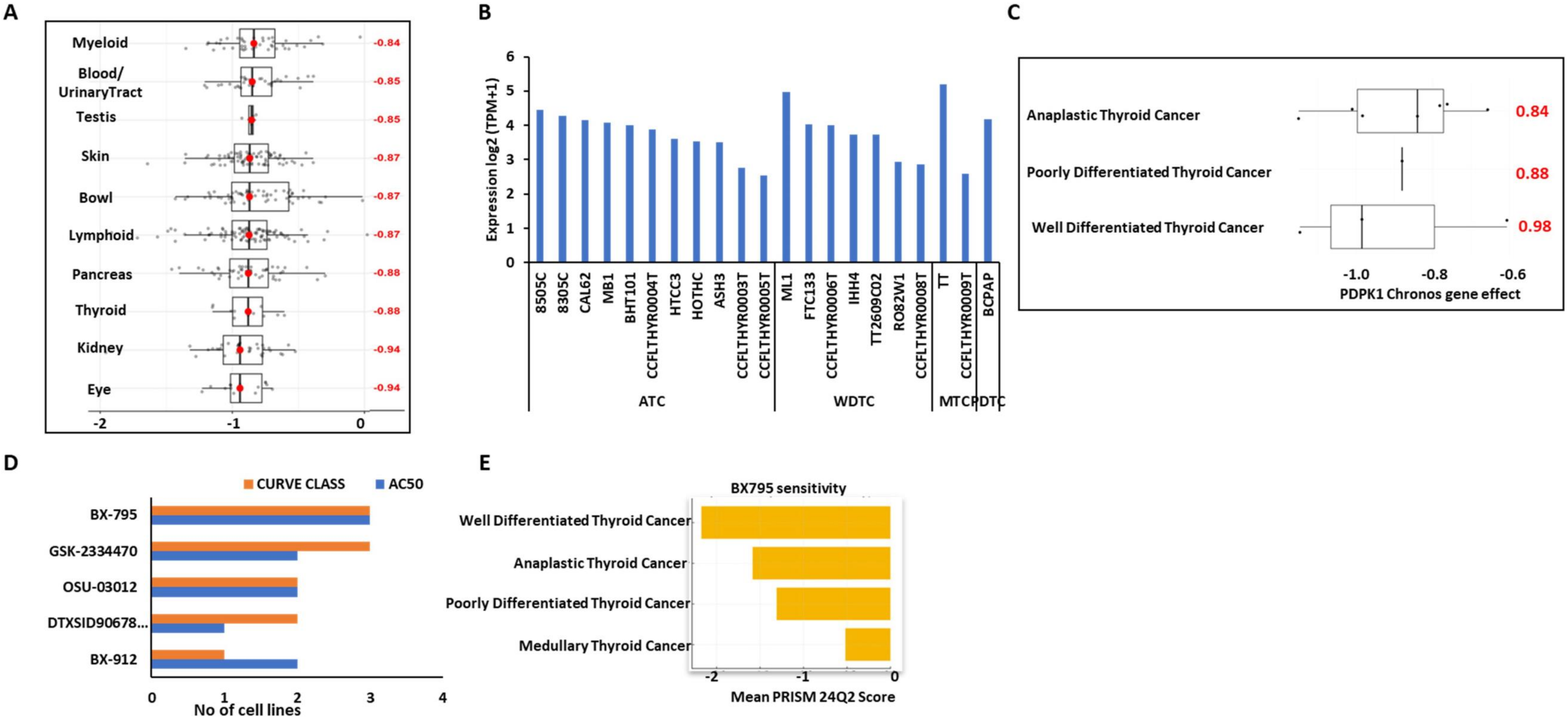

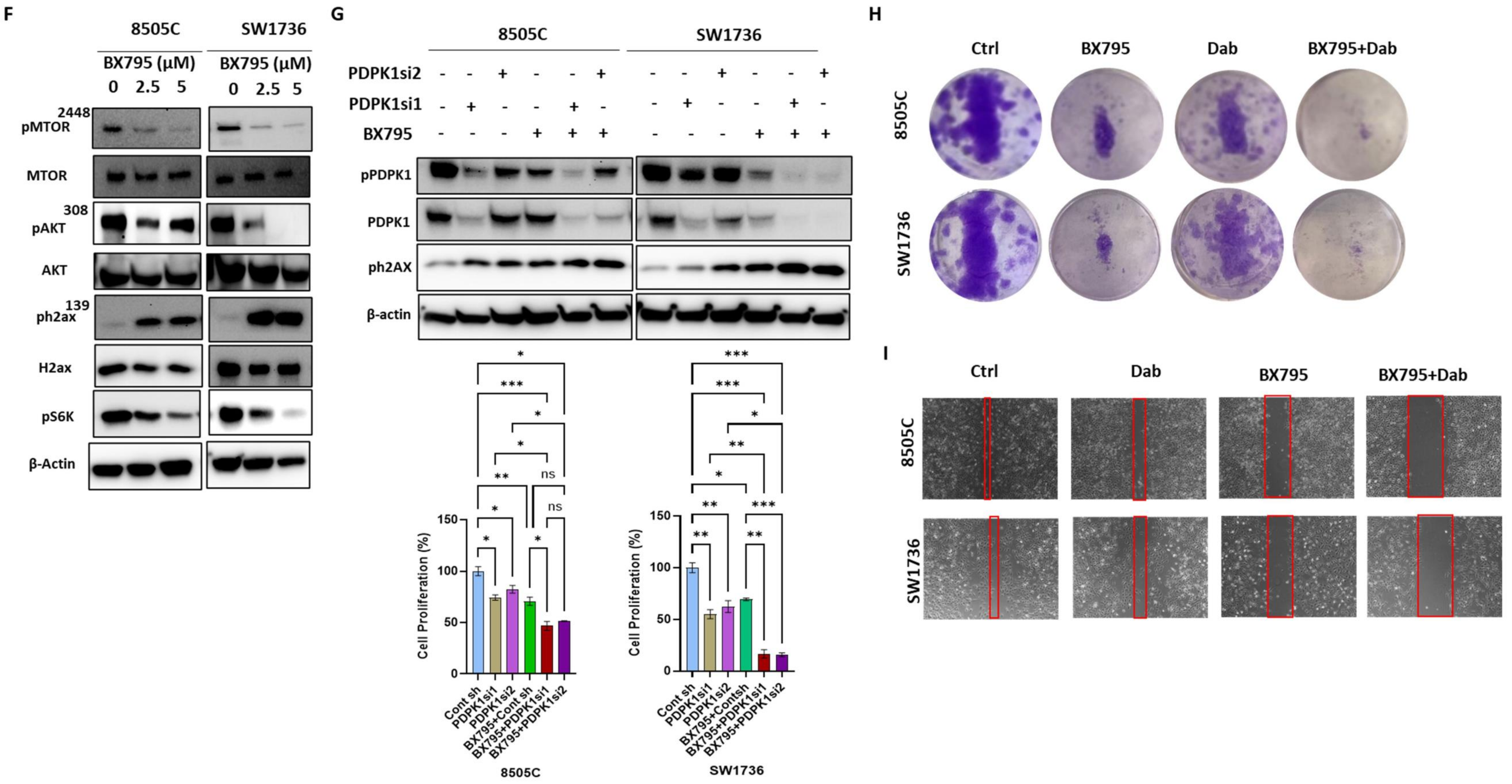

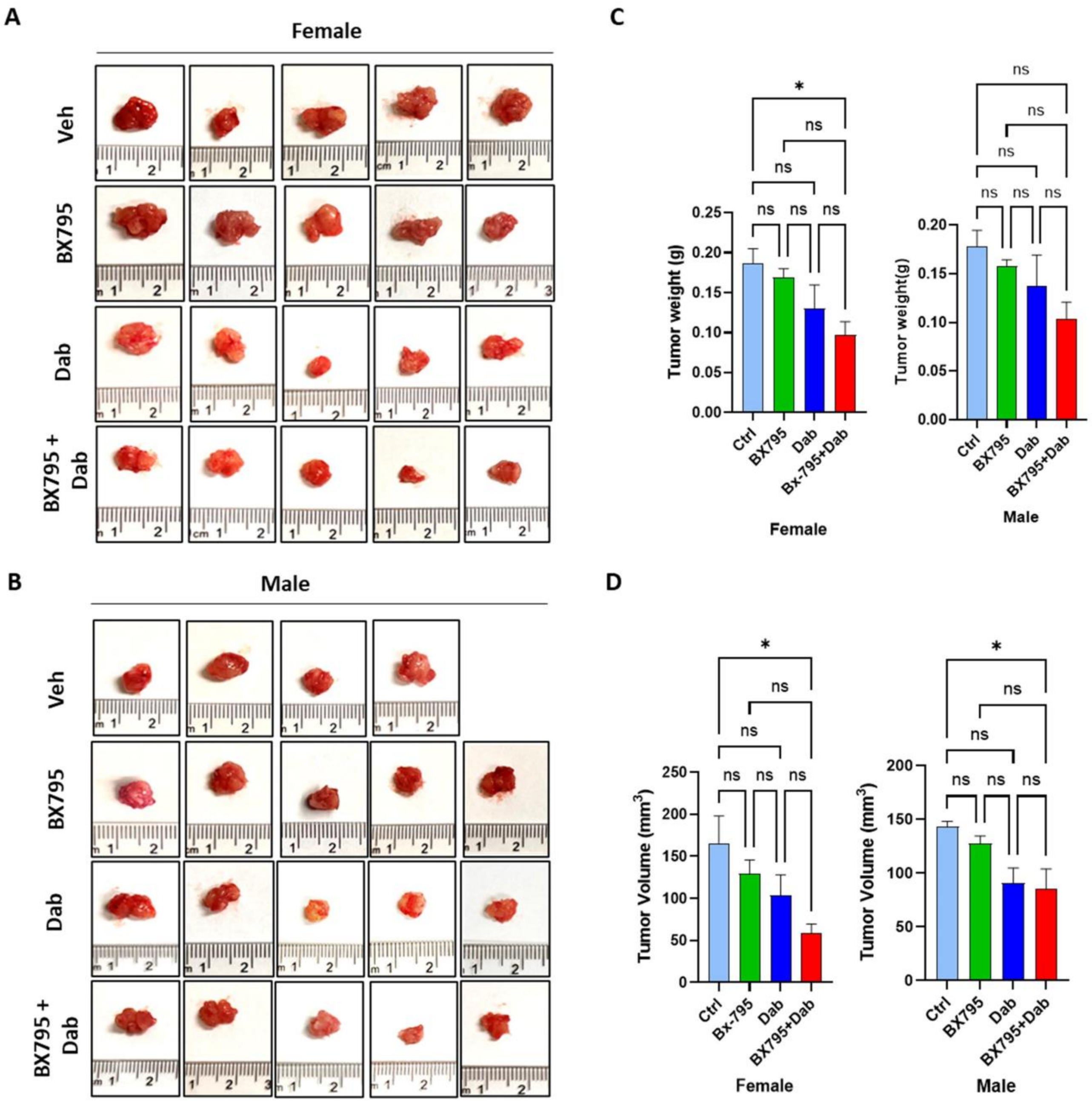

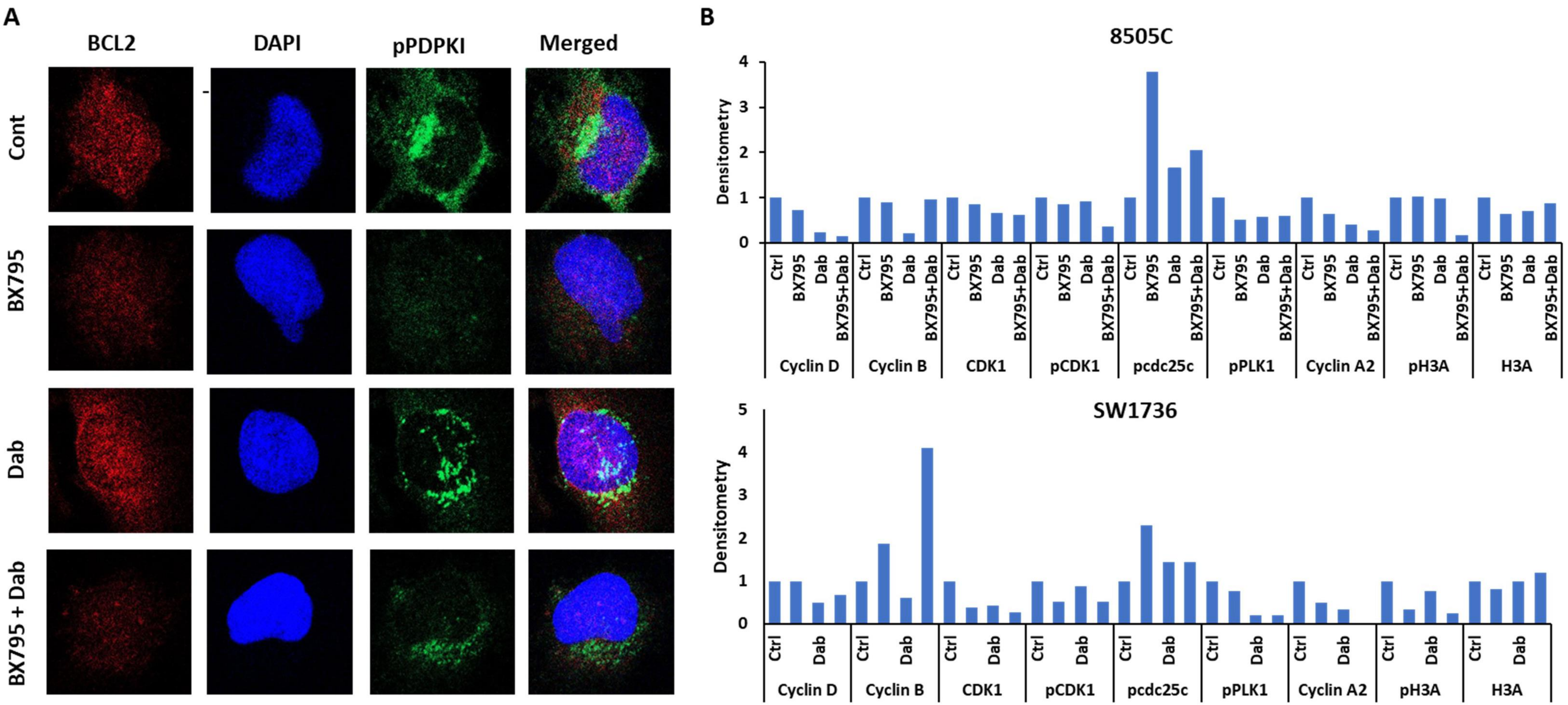

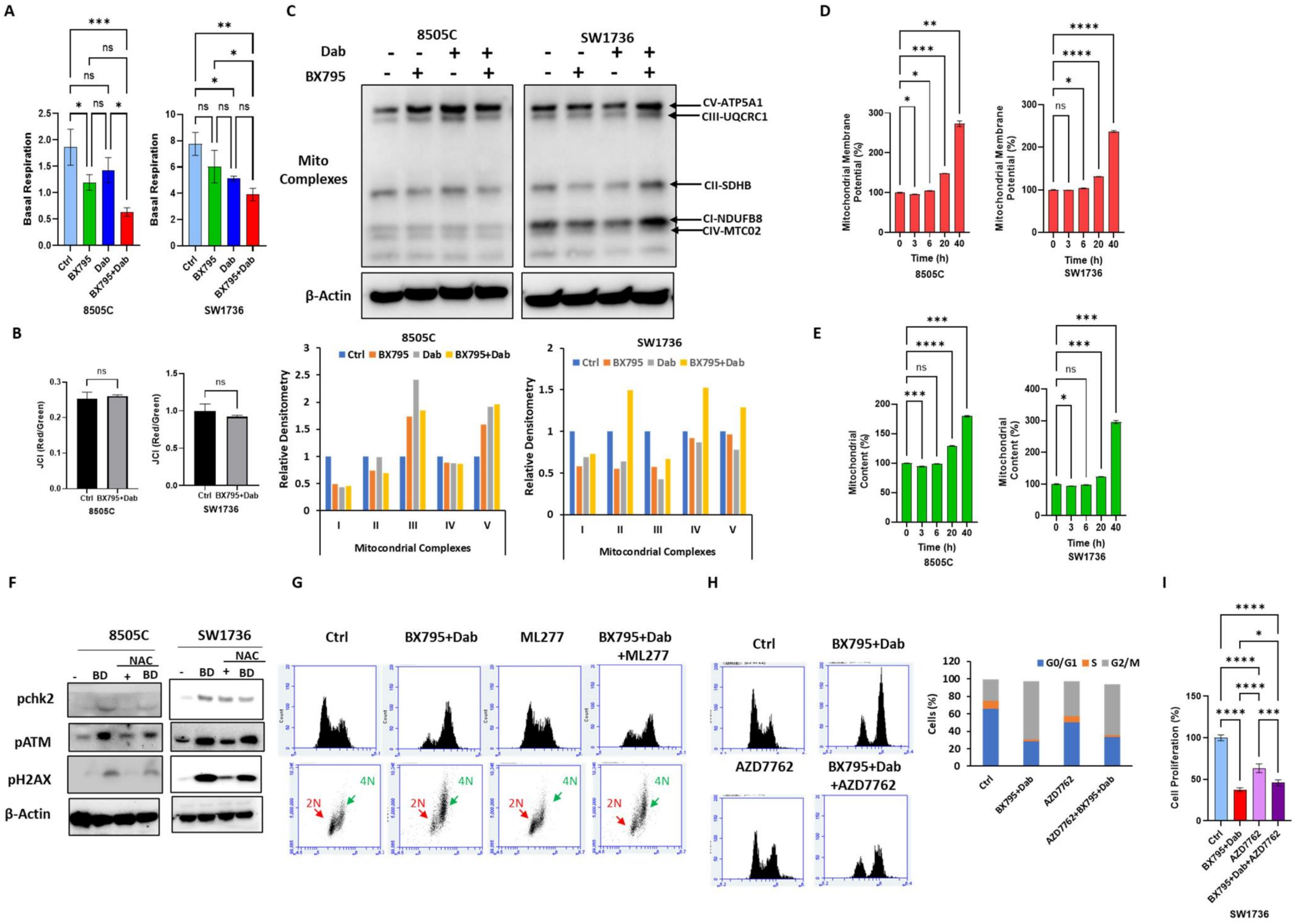
